# Structural and systems characterization of phosphorylation on metabolic enzymes identifies sex-specific metabolic reprogramming in obesity

**DOI:** 10.1101/2024.08.28.609894

**Authors:** Tigist Y Tamir, Shreya Chaudhary, Annie X Li, Sonia E Trojan, Cameron T Flower, Paula Vo, Yufei Cui, Jeffrey C Davis, Rachit S Mukkamala, Francesca N Venditti, Alissandra L Hillis, Alex Toker, Matthew G Vander Heiden, Jessica B Spinelli, Norman J Kennedy, Roger J Davis, Forest M White

## Abstract

Coordination of adaptive metabolism through cellular signaling networks and metabolic response is essential for balanced flow of energy and homeostasis. Post-translational modifications such as phosphorylation offer a rapid, efficient, and dynamic mechanism to regulate metabolic networks. Although numerous phosphorylation sites have been identified on metabolic enzymes, much remains unknown about their contribution to enzyme function and systemic metabolism. In this study, we stratify phosphorylation sites on metabolic enzymes based on their location with respect to functional and dimerization domains. Our analysis reveals that the majority of published phosphosites are on oxidoreductases, with particular enrichment of phosphotyrosine (pY) sites in proximity to binding domains for substrates, cofactors, active sites, or dimer interfaces. We identify phosphosites altered in obesity using a high fat diet (HFD) induced obesity model coupled to multiomics, and interrogate the functional impact of pY on hepatic metabolism. HFD induced dysregulation of redox homeostasis and reductive metabolism at the phosphoproteome and metabolome level in a sex-specific manner, which was reversed by supplementing with the antioxidant butylated hydroxyanisole (BHA). Partial least squares regression (PLSR) analysis identified pY sites that predict HFD or BHA induced changes of redox metabolites. We characterize predictive pY sites on glutathione S-transferase pi 1 (GSTP1), isocitrate dehydrogenase 1 (IDH1), and uridine monophosphate synthase (UMPS) using CRISPRi-rescue and stable isotope tracing. Our analysis revealed that sites on GSTP1 and UMPS inhibit enzyme activity while the pY site on IDH1 induces activity to promote reductive carboxylation. Overall, our approach provides insight into the convergence points where cellular signaling fine-tunes metabolism.

**Summary Statement:** By employing a multi-disciplinary approach we stratify structural features of phosphorylation sites on metabolic enzymes, map the systems level changes induced by obesity, identify key pathways with sex specific phosphoproteomic responses, and validate the functional role of phosphorylation sites for select enzymes.

## Introduction

Throughout evolution, the battle for survival has been paved by how the cell adapts to environmental stressors by rewiring signaling and metabolic networks. Coordination of metabolic networks is tightly regulated from sensing fluctuation in metabolite levels to modulation of enzyme activity and expression. There are many layers of context specific control systems that are optimized for robust metabolic output; post-translational modifications (PTMs) on metabolic enzymes represent one such layer^1,2^. While expression of metabolic enzymes sets the stage for potential pathway activity, adaptive response and regulation are known to stem from transient events like PTMs that can regulate responses on the seconds to minutes timescale^3–6^. Protein phosphorylation on serine (S), threonine (T), and tyrosine (Y) residues is one of the most prevalent PTM^1,7,8^. Phosphorylation-mediated signaling regulation of metabolism is an important component of network level modulation of bioenergetics as well as homeostasis. Both dynamic and chronic perturbations alter phosphorylation of metabolic enzymes on highly conserved residues^6,9–12^. A wide range of signaling and metabolic rewiring governed by phosphorylation has been reported during adaptive response to metabolic stressors (e.g., chemotherapy drugs, inflammation, and high fat diet)^13–16^. As a result, dysregulation of both signaling and metabolism are commonly observed in disease states such as obesity and cancer^11,13,17–19^.

One of the most important processes regulating the adaptive response to metabolic stressors is the Oxidative Stress Response (OSR) pathway, which prevents oxidative damage via a complex interconnected set of enzymatic and non-enzymatic scavengers that maintain reactive oxygen/nitrogen/carbonyl species (ROS/RNS/RCS) at a tolerable level^20–24^. Dysregulation of the OSR due to overactive or deficient antioxidant capacity can cause damage to biomolecules such as DNA, RNA, protein, and lipids^23,25–27^. As a result, OSR dysregulation is a hallmark of many human diseases including cancer, inflammation, neurodegeneration, cardiovascular diseases, obesity, diabetes, and ageing^13,24,28–30^. Enzymes that govern OSR, which are often altered in diseases, are tightly regulated at the transcriptional, translational, and post-translational levels^23,24,29,31,32^. These enzymes prevent damage from oxidative stress related internal (e.g., metabolic waste) and external (e.g., xenobiotics) cellular insults^27,29^. A number of phosphorylation events have been identified on antioxidant enzymes, but only a limited set have been well characterized in a perturbation and tissue specific manner ^8,9,11,33–39^. It is paramount that we understand the intricate regulatory process of the OSR metabolon, and how antioxidant enzymes integrate signaling into rapid response can elucidate not only the process of life but also how it is altered in diseases.

Advances in large scale phosphoproteomics and enrichment techniques have improved our capacity to identify an increasing number of phosphorylation sites on metabolic enzymes. While some of these sites are functionally characterized, a systemwide analysis of the phosphorylation-modulated metabolic network remains largely unexplored. Moreover, there exist comprehensive genomic, epigenetic, transcriptomic, proteomic, and metabolite level annotations of metabolic networks, yet there are limited systems level analyses of how phosphorylation events on metabolic enzymes contribute to overall metabolism and network topology. In order to understand the many ways metabolic rewiring can be shaped by perturbations, we need to evaluate the combined impact of phosphorylation events on multiple metabolic enzymes and their concerted effect on the system.

Here we address this challenge by taking a systems approach that integrates computational structural analysis, phosphoproteomics, metabolomics, and biochemical validation of phosphorylation mediated regulation of metabolic enzymes. We specifically interrogate pY on metabolic enzymes using a comprehensive analysis of an *in vivo* model of high fat diet (HFD)- induced obesity where we map out metabolic tuning that alters redox homeostasis. Furthermore, we employ a targeted mass spectrometry-based approach to interrogate pathway specific phosphorylation events and apply computational modeling to identify pY sites associated with altered metabolic output. Lastly, we validate model predictions by measuring the functional role of selected pY sites using a CRISPRi-rescue system with phosphomimic or phosphodeficient enzyme variants in biochemical and isotope tracing metabolomics analysis. We demonstrate that pY sites in functional domains and dimerization interfaces provide a concerted additive coordination of metabolic pathways to direct dynamic changes in chronic metabolic reprogramming of obesity.

## Results

### The published metabolic phosphoproteome is enriched for oxidoreductases

To evaluate the role of phosphorylation sites on metabolic enzymes we curated the published human phosphoproteome dataset available through PhosphoSitePlus (PSP)^40^, filtering phosphorylation sites on serine (S), threonine (T), and tyrosine (Y) residues to focus on metabolic enzymes as annotated by the Kyoto Encyclopedia of Genes and Genomes (KEGG) metabolic pathway (Figure 1A, Table S1)^41,42^. Of the 239885 sites in the dataset, 57589 were on proteins classified as enzymes with 13679 of these sites occurring on metabolic enzymes of which 877 enzymes have published structural data. Phosphorylation sites on metabolic enzymes comprise 6% of the measured phosphoproteome, however of the 1599 metabolic enzymes we curated from KEGG, 1353 of them have previously reported phosphosites. Thus, nearly 85% of metabolic enzymes in the human proteome are phosphorylated (Figure 1B). Of these phosphosites on metabolic enzymes, 50% are phosphoserine (pS), 25% are phosphothreonine (pT), and 24% are phosphotyrosine (pY), respectively. While overall cellular pY stoichiometry is of low abundance, the current set of published phosphoproteomic studies favor contexts that enrich for pathways that induce pY (e.g., altered growth factor/nutrient conditions). Gene level pathway enrichment analysis of the published metabolic enzyme phosphoproteome revealed overrepresentation of enzymes in lipid metabolism, carbohydrate metabolism, metabolism of nucleotides, fatty acid and phospholipid metabolism, and regulation of oxidation (Figure 1C, Table S1)^43–45^. The most significantly enriched molecular functions of phosphorylated enzymes were oxidoreductase activity, especially enzymes that bind NAD/NADP (Figure 1D, Table S1)^43,44^. These results suggest that phosphorylation is a common PTM on metabolic enzymes, and that the measured metabolic enzyme phosphoproteome is enriched for enzymes that generate, regulate, and resolve oxidative stress.

**Figure 1.**
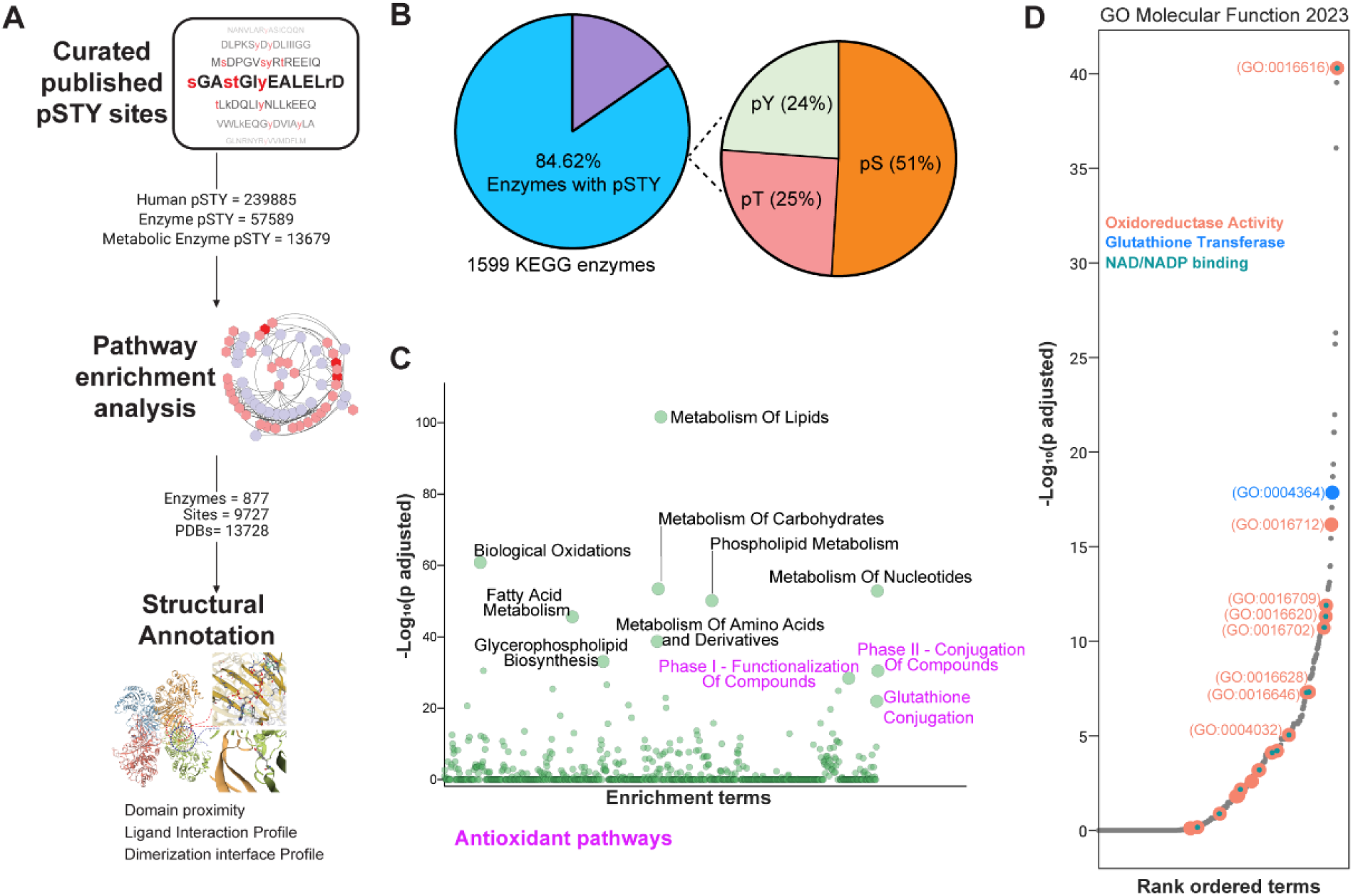
Oxidative pathways and oxidoreductases are overrepresented in the published metabolic enzyme phosphoproteome. (A) Workflow depicting structural annotation of phoshosites on PDB structures. **(B)** Pie chart representing that the majority of metabolic enzymes are phosphorylated as annotated by cross-referencing proteins curated from KEGG and published phosphosites in PSP. **(C)** Gene level pathway enrichment analysis of phosphorylated metabolic enzymes performed using Reactome gene set and Enrichr analysis tool. Data is represented as Manhattan plot for enrichment terms and p-values from Fisher’s exact test was adjusted by Benjamini-Hochberg procedure. **(D)** Molecular function enrichment analysis using GO Molecular Function gene set and Enrichr plotted by ranked GO terms illustrating that NAD/NADP cycling oxidoreductases are overrepresented in the published metabolic enzyme phosphoproteome, and Fisher’s exact test p-values were adjusted by Benjamini-Hochberg procedure.

### Structural annotation of phosphosites on metabolic enzymes reveals pY is overrepresented in functional domains and dimerization interfaces

Given the prevalence of phosphorylation sites on metabolic enzymes, we next asked if we can infer the functional role of these sites based on structural features. To characterize the structural features of phosphosites on metabolic enzymes, we focused on enzymes with published structural data available through the Protein Data Bank (PDB)^46^. We used structures with resolution < 2.6Å where side chains are well resolved, and Rfree – Rwork < 0.05 to filter for crystal structures with refinement cross-validation that aligns with the experimental data, thus only selecting high quality structures (Figure 2A, Table S2)^47^. Unfortunately, there are very limited structures containing phosphorylation sites due to the transient nature of this PTM, therefore we obtained structural information from the unphosphorylated (apo) residues as a proxy to evaluate the structure-function relationship of phosphosites on metabolic enzymes. To measure the proximity of each phosphosite to enzyme functional domains, we first curated annotated domains from Uniprot-KB, and mapped them onto PDB structures^48^. Using PyMol, we measured the distance from the hydroxyl group of the apo residues to the center of mass (COM) of any defined functional domains of the enzyme, then averaged the distances across all available PDB structures for each enzyme. Of the published phosphosites, 50% were approximately 23Å from the COM of functional domains (Figure 2B, Table S3), suggesting that upon phosphorylation these residues are likely to influence domain structure and affect enzyme function directly, especially given the size and charge of the phosphate moiety. Moreover, phosphorylation sites were more frequently observed in substrate binding, cofactor binding sites, and active sites, respectively (Figure 2C, Table S3). We then asked which phosphorylated residues were most enriched within 23Å of each domain type. Overall, substrate binding and active sites had the greatest enrichment for phosphorylated residues (Figure 2D, Table S4). Intriguingly, despite low cellular stoichiometry, pY was most commonly overrepresented across substrate binding, active sites, and cofactor binding sites.

**Figure 2.**
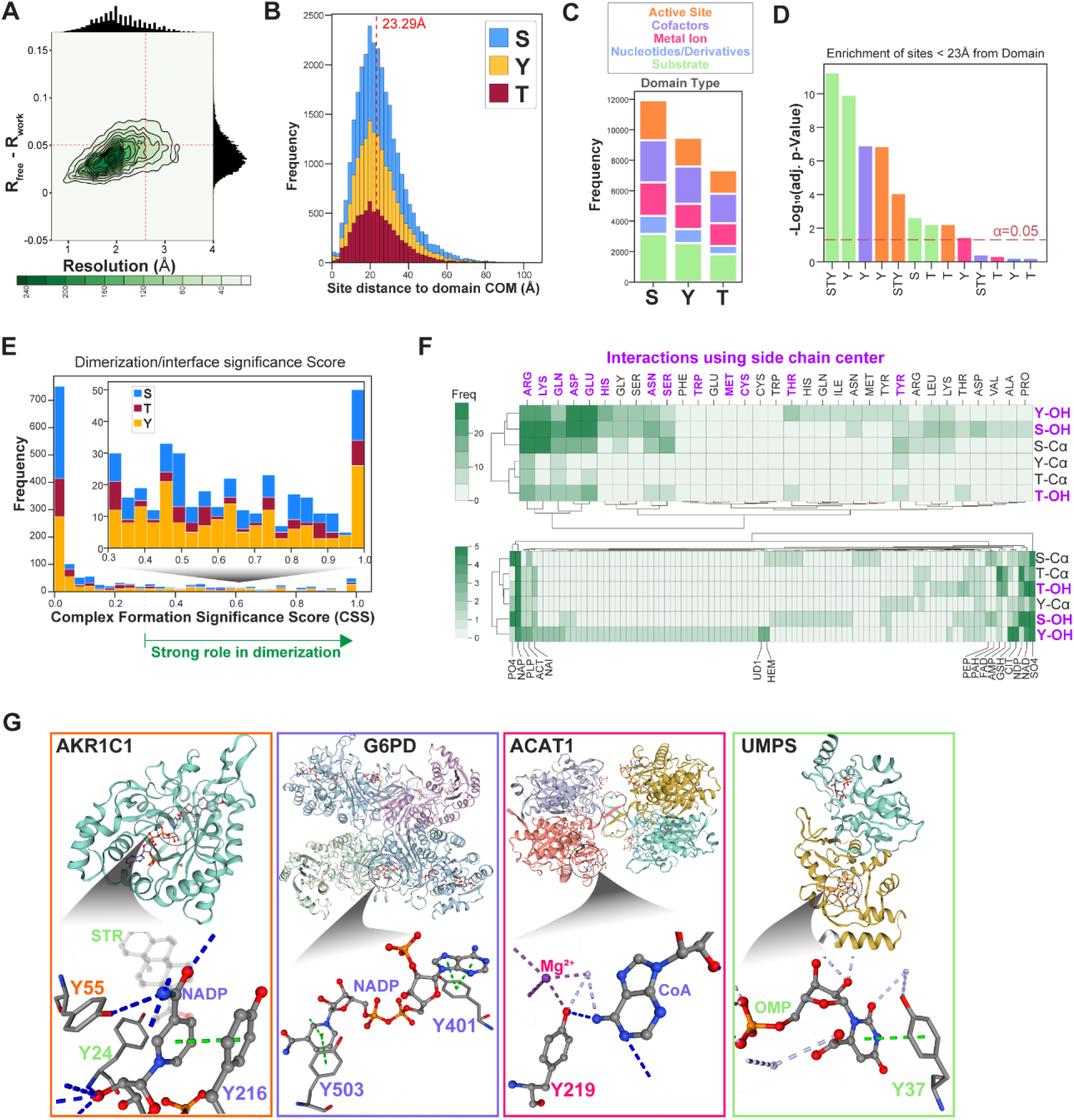
Structural annotation of phosphosites on metabolic enzymes identifies overrepresentation of pY in functional and dimerization domains. (A) Joint plot representing assessment of published structural data for metabolic enzymes based on resolution and refinement cross-validation. High quality structures are indicated by red dotted lines at resolution ≤ 2.6Å and Rfree – Rwork ≤ 0.05 to filter for high quality x-ray crystal structures. **(B)** Stacked histogram plot of phosphosite distance from domains. Protein functional domains curated from UniprotKB were annotated on corresponding structures using PyMol. For each phosphosite on a given enzyme, distance was measured from the hydroxyl group of the apo residue to the center of mass (COM) of any defined functional domain, distances were averaged across all available PDB structures of each enzyme for phosphosite-domain pairs. Stacked histogram plot represents frequency of distances for each pair of phosphosite and functional domain, where 50% of the data is indicated by red dotted line at 23.29Å. **(C)** Functional domains for each enzyme were classified into domain types based on UniprotKB annotation, and stacked bar plot represents the proportion of phosphosite residues within 23Å of each domain type. **(D)** Bar plot representation of hypergeometric-distribution test with multiple hypothesis correction of the cumulative distribution function done via Benjamini-Hochberg test. (**E)** Stacked histogram plot of complex formation score for dimer interfaces that contain phosphosites. High quality protein structures of metabolic enzymes were used to identify dimerization interfaces and the residues that form contacts important for complex formation using PISA analysis tool. The data was filtered for phosphosites only, and stacked bar plot represents frequency of complex formation score (CSS) obtained from PISA. CSS≥0.3 indicates that the interface plays a strong role in dimerization. **(F)** Heatmap for the frequency of interaction between phosphosite hydroxyl group (OH) or backbone (Ca) with residues and ligands found in the interface region. **(G)** 4 examples of phosphosites on metabolic enzymes illustrate the versatile role phosphosites serve in functional domains. Structure of AKR1C1 showing hydrophobic interactions (blue dotted line) of Y55 with cofactors (NADH or NADPH), Y24 with steroid substrates (progesterone shown here), and pi-stacking interaction (green dotted line) of Y216 with the nicotinamide ring of NADP+. G6PD protein structure showing pi-stacking interactions between Y401, Y503, and NADP within the interface of two monomers. ACAT1 structure showing interaction between the hydroxyl group on Y219 with cations (e.g., K+, Mg2+) as well as a water bridge and hydrogen bond with the adenosine ring of Coenzyme A (CoA). Structure of UMPS showing hydrogen bonds and pi-stacking interaction between Y37 and OMP. Illustrations were generated using SwissModel^103,104,138^ and PLIP^105^ tool and the following PDBs: AKR1C1 (1mrq)^139^, ACAT1 (2iby)^140^, G6PD (7sni)^55^, UMPS (2wns)^102^.

While curating functional domains from Uniprot allowed for annotation of domain proximity, it was limited in characterizing dimerization domains which are not commonly defined in the database. Therefore, we utilized the Proteins, Interfaces, Structures and Assemblies (PISA) tool from PDBe to obtain the properties of residues within multimeric interfaces^49–51^. To evaluate whether phosphorylated interface residues play a significant role in dimerization we obtained the interface complex-formation significance score (CSS), which measures how much the interface contributes to intramolecular interaction (i.e., dimerization). While the majority of interfaces containing phosphosites had a low CSS, those with CSS ≥ 0.3 were most commonly pY and pS, respectively (Figure 2E, Table S5). This suggests that pY and pS in interface regions of multimeric enzymes most likely alter dimerization. To ascertain the role of the OH atoms on phosphosites of interface residues, we identified common interaction partners of interface phosphosites. We grouped phosphosites into two groups based on which atoms form interactions: side chain centers (R groups of residues where PTMs are commonly added) or protein backbone (C-alpha, Cα). The most common interactions of interface phosphosites happen through the phosphate accepting hydroxyl group, especially for pY and pS. While pS can form interactions with both the hydroxyl group and back bone atoms, pY is unique in that most, if not all, interactions take place through the hydroxyl group. Furthermore, interface pY and pS in their apo form interact most commonly with the side chains of charged and polar amino acids such as Arginine (ARG), Lysine (LYS), Glutamine (GLN), Aspartate (ASP), Glutamate (GLU), and Histidine (HIS), respectively (Figure 2F, Table S5). Similarly, for interface regions with small molecule ligands, pY and pS show the most interaction with NAD, NADP, NADPH, GSH, and SO4. Although there were some structures with free phosphate groups (PO4), only hydroxyl groups from pS sites were observed as interactors. In general, interface interactions have an average distance of < 3Å (Figure S1A, Table S5), suggesting that the modification of the phosphosite hydroxyl with a ∼4 - 5Å negatively charged phosphate moiety will have major ramifications for the dimerization and function of multimeric enzymes. Therefore, our computational structural analysis underscores that phosphosites on metabolic enzymes are within close proximity of functional and dimerization domains, and that pY sites, in particular, present unique structural features that highly correlate structure to function.

To illustrate the many structural characteristics of phosphosites, particularly for pY, we selected examples of sites on metabolic enzymes, including previously characterized and uncharacterized sites on AKR1C1, G6PD, ACAT1, and UMPS (Figure 2G). AKR1C1 is a cytosolic enzyme that catalyzes reduction of steroids and facilitates lipid metabolism, where pY sites Y24, Y55, and Y216, are localized in the substrate, active, and cofactor binding sites, respectively ^52,53^, although functional characterization has yet to be determined. Sequence alignment of all 14 enzymes in the AKR family shows that all 3 pY sites are highly evolutionarily conserved: Y55 is in all 14, Y216 in 11, and Y24 is in 6 (Figure S1B). G6PD is another well-known modulator of cellular redox homeostasis via the pentose phosphate pathway (PPP), and is a target of both growth factor signaling and direct substrate of SRC family kinases (SFK)^35,54–56^. Phosphorylation at Y401 and Y503 have been characterized as inducers of enzyme activity by increasing the rate of NADP+ cycling. ACAT1 is involved in the anabolism and catabolism of ketone bodies in the mitochondria during fatty acid oxidation^57^. The functional role of ACAT1 Y219 phosphorylation is undefined, where the hydroxyl group interacts with cations and water suggesting that phosphorylation of Y219 would alter ACAT1 activity. Of note, Y407 on ACAT1 (not shown here) has been characterized as regulator of multimeric enzyme complex formation^58^. Lastly, UMPS catalyzes the last step of *de novo* pyrimidine syntheses by producing UMP from orotate^59–62^. Residue Y37 forms a pi-stacking interaction and a water bridge with the orotate ring of OMP. While highly reported in large scale phosphoproteomics analysis, pY37 on UMPS is not characterized. These 4 examples of the apo resides from high quality structures suggest that pY sites in these enzymes likely alter the structural properties of functional domains, affect enzyme activity, and thereby alter flux through associated metabolic networks. Overall, structural annotation of phosphorylation sites on metabolic enzymes elucidated that while pY is rare in the cellular phosphoproteome, it is enriched in functional domains with potential consequence for enzyme activity as well as systemic metabolism.

### High fat diet induced change in metabolites are associated with redox metabolism

While our structural characterization highlighted the prevalence of pY sites associated with regulatory domains of metabolic enzymes and that many of these sites would impact enzymatic activity and metabolic pathways, we wanted to assess whether these sites could have regulatory function in the context of metabolic stressors. To address this question, we employed a diet induced obesity (DIO) mouse model with integrative phosphoproteomic and metabolomic analyses of mouse livers to characterize the response of pY-mediated signaling networks, fatty acid metabolism and biological oxidation pathways to high fat diet (HFD). DIO sensitive male and female C57BL/6J mice given normal chow diet (NCD) or HFD livers were collected for multiomic analysis (Figure 3A, Table S6). To quantify the physiological impact of HFD induced changes, we assessed body weight, fat/lean mass, blood glucose, and insulin levels. Compared to NCD, both male and female mice on HFD gained total body mass as well as fat mass, but not lean mass (Figure 3B, S2A, Table S6). While the fold change of average body weight gained in HFD compared to NCD was similar between sexes (males ≈ 1.6, females ≈ 1.5), the fold change of fat mass gained by males (HFD/NCD ≈ 5.1) was higher than females (HFD/NCD ≈ 3.8). Fasted blood glucose (FBG) levels were increased in both sexes with males having higher fold change when HFD was normalized to NCD (males ≈ 2.3, females ≈ 1.9). In contrast, blood insulin levels were only significantly elevated by HFD in males (p=0.0057) but not females (p=0.1475) (Figures 3C, 3D, Table S6). Taken together, and in line with the literature, there were significant physiological changes in both males and females indicative of obesity, however only males exhibited metabolic syndrome/insulin resistance^63–65^.

**Figure 3.**
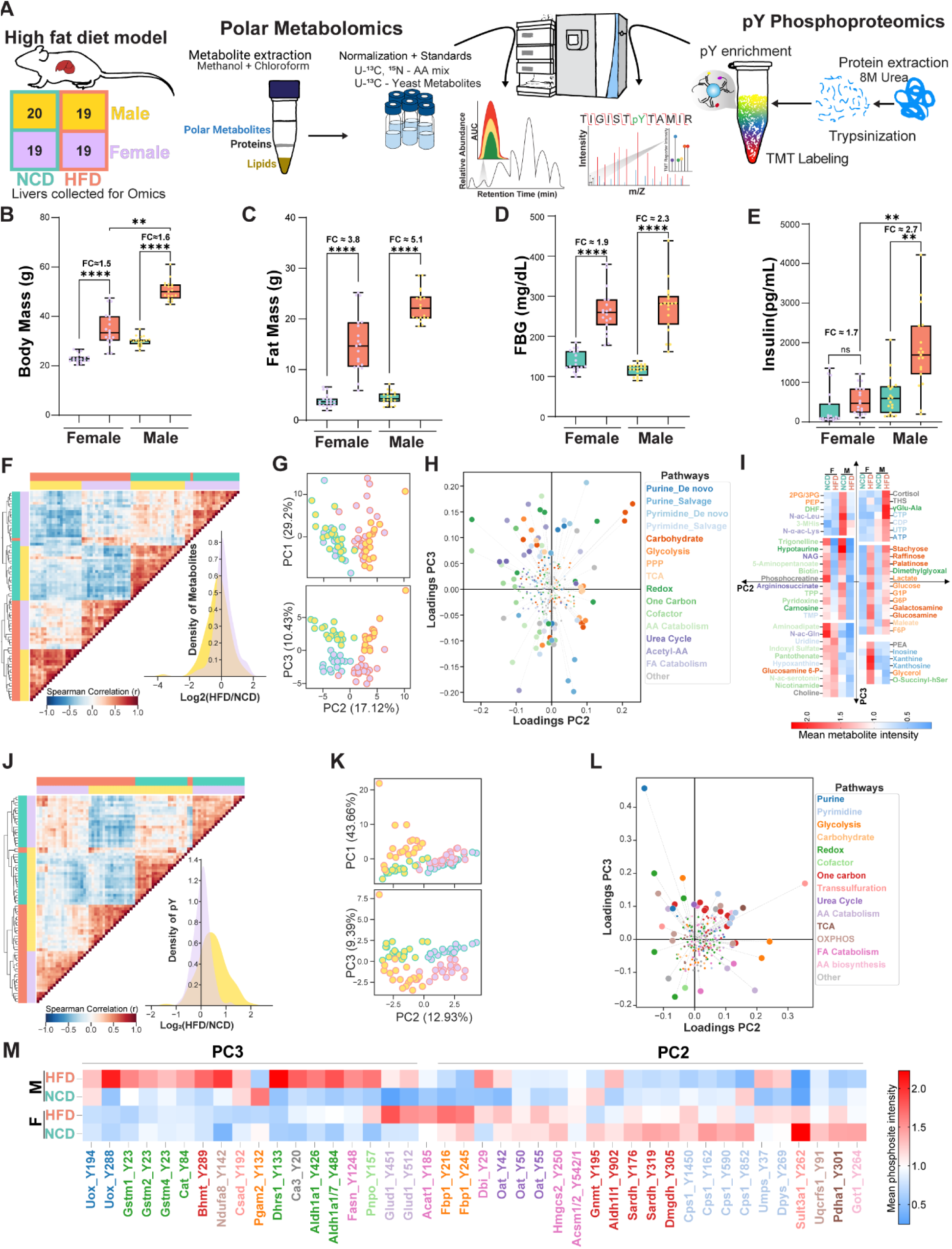
HFD induced obesity results in sex-specific phenotypic, phosphoproteomic, and metabolic changes. (A) Schematic of *in vivo* experiment in C57BL6/J mice fed high fat diet (HFD, 60% kcal from fat, N=19) or normal chow diet (NCD, 30% kcal from fat, N=19) for 16 weeks, and dry pulverized livers were collected for multiomic analysis of steady state polar metabolites and pY phosphoproteome. **(B-E)** Physiological measurements of total body mass (g), fat mass (g), fasted blood glucose (FBG, mg/dL), and blood insulin (pg/mL) levels. Data points represent individual animals. Two-way ANOVA with multiple comparison (Kruskal-Wallis test) was performed comparing corresponding NCD with HFD groups and statistical significance was assigned as follows: * p < 0.05, ** p < 0.01, **** p < 0.001, ns = not statistically significant. **(F)** Heatmap of spearman correlation analysis of polar metabolomics data with hierarchical clustering. KDE plot for the distribution of metabolite fold changes represented as Log2(HFD/NCD) demonstrating decreased metabolites in males evidenced by left-shift in the distribution. **(G)** Scatter plot of PCA on polar metabolite measurements, data points represent individual animals. **(H)** Scatter plot of metabolites that drive PCA clustering (loadings) along PC2 and PC3 color-coded by pathway. Data points represent individual metabolites. **(I)** Heatmap representation of metabolites differentially regulated by diet and sex in PC2 and PC3. **(J)** heatmap representation of spearman correlation analysis and hierarchical clustering of targeted pY phosphoproteomics analysis on metabolic enzymes in response to HFD. KDE plot showing distribution of pY site Log2(HFD/NCD) demonstrating an increase (right-shift) in males but not females. **(K)** Scatter plot representation of PCA on pY sites along PC1, PC2 and PC3. Data points represent individual animals. **(L)** Loadings scatter plot of pY sites that drive clustering across PC2 and PC3, color coded according to metabolic pathway of each enzyme. Data points represent pY sites. **(M)** Mean intensity of pY site drivers of PC2 and PC3 clustering represented as heatmap across sex and diet.

To determine altered metabolites in response to HFD, we performed mass spectrometry based steady state polar metabolomics analysis. We identified and quantified 260 polar metabolites across 77 samples. Spearman correlation analysis and hierarchical clustering show that HFD induced changes in metabolites are anticorrelated to NCD and that there is a distinct metabolic signature between males and females (Figure 3F). In line with phenotypic observations, males exhibited a greater response to HFD compared to females, with a large number of metabolites being decreased in HFD compared to NCD fed male mice (Figure 3F, Table S7). Principal Component Analysis (PCA) revealed that the majority of variance across diet and sex were captured on PC2 (17.12%) and PC3 (10.43%), respectively (Figure 3G). The loadings, drivers of differential clustering, for males on HFD were nucleotides from *de novo* purine/pyrimidine biosynthesis pathways, while those on NCD were differentiated by metabolites from urea cycle, fatty acid (FA) catabolism, and N-acetyl amino acids (Figure 3H, *S*2E). For example, inosine, xanthine, and xanthosine were all elevated by HFD in females but not males, while nucleotides from pyrimidine synthesis and cortisol were only increased in males (Figure 3I). There is strong evidence that inosine increases whole-body energy homeostasis which prevents obesity by reducing lipid deposition and increasing adipose tissue thermogenesis^66–70^. In contrast, increased cortisol or stress hormone levels are strongly associated with adiposity, thus high cortisol and tetrahydroxycortisol (THS) levels in males on HFD (Figure 3I) is in line with observations from human trials^71^. Our data suggest that females exhibit less severe metabolic syndrome in part due to capacity to modulate insulin sensitivity and whole-body bioenergetics through a combination of nucleotide metabolism as well as redox homeostasis.

### Targeted phosphoproteomics reveals that HFD results in sexually dimorphic changes of pY on metabolic enzymes

In parallel with metabolomic analysis, we performed untargeted pY phosphoproteomics on 77 samples, leading to identification of 589 pY sites, including 174 pY sites on metabolic enzymes quantified in all samples. Spearman correlation analysis shows anticorrelation of pY sites altered due to both diet and sex (Figure S2B). PCA of untargeted pY analysis shows that variance captured by PC1 (24.6%) explains differences between NCD and HFD but only in male livers (Figure S2C), whereas PC2 (12.07%) and PC3 (8.96%) capture variance across both sex and diet. Sites that contribute to PC1 include activating pY sites on signaling proteins downstream of insulin and growth factor signaling (e.g., Mapk1/3_Y185, Y205) (Figure S2D).

To get deeper coverage of phosphosites and overcome the limit of detection for pY sites on metabolic enzymes we employed targeted phosphoproteomics. We generated an inclusion list of all reported sites on metabolic enzymes from PSP and our untargeted analysis, and filtered for sites on enzymes in pathways represented in our polar metabolomics dataset. We captured 269 pY sites on metabolic enzymes across 77 samples, and measured over 1.5-fold more pY sites on metabolic enzymes compared to our untargeted analysis. Targeted pY phosphoproteomics recapitulated the anticorrelation of pY signatures for both diet and sex (Figure 3J). In agreement with the metabolite and phenotypic data, signaling networks in male mice were more strongly affected by HFD compared to female mice (Figures 3J, S2F, Table S5). PCA of targeted pY analysis effectively clustered samples by diet and sex in PC2 (12.93%) and PC3 (9.39%), respectively (Figure 3K). The variance captured by PC2 was observed for pY sites on metabolic enzymes of one carbon, pyrimidine, and redox metabolism (Figures 3L, 3M). Meanwhile, the variance captured by PC3 is directed by pY sites on enzymes of purine metabolism, fatty acid (FA) catabolism, and oxidative phosphorylation (OXPHOS) (Figures 3L, 3M).

To visualize the network level changes in both omics’ datasets, we constructed a metabolic pathway map (Figure S3) representing metabolites and pY sites associated enzymes. Enzymes involved in glutamine/glutamate biosynthesis (e.g., Glud1, Glul), redox homeostasis (e.g., Aldh1a1/7, Rida), and glycolysis (e.g., Pgam1) were hyperphosphorylated in response to HFD (Figures 3M, S3). However, enzymes regulating one carbon metabolism (e.g., Gnmt, Sardh) and urea cycle (e.g., Cps1, Ass1) had decreased phosphorylation following HFD. Since there was a difference in severity of HFD induced physiological changes between males and females, we asked which major pathways were most differentially altered. We queried the structural annotations of pY sites to elucidate their potential functional role. In line with our interrogation of the published phosphoproteome, pY sites on redox enzymes were most prevalent in functional domains (Figure 4A). Males exhibited increased pY on redox enzymes in glutathione biosynthesis (GSS, GSR, GSTs), quinone reduction (Cryz, Coq5), and aldehyde detoxification (Aldh1a1/7). Among the redox enzymes, phosphorylation of GSS Y270 is the only characterized site, shown to inhibit enzyme function by creating steric hindrance of substrate binding (i.e., Ɣ-GluCys) ^72–74^. Additionally, pY on enzymes in fatty acid metabolism and cataplerosis pathways from acetyl-CoA were higher in males than females with the majority of sites being in enzyme interface regions (Figure 4A). Together these data suggest that the phenotypic outcomes of HFD-induced obesity in male mice are associated with altered redox homeostasis governed by signaling modulation of metabolic enzymes and their function.

**Figure 4.**
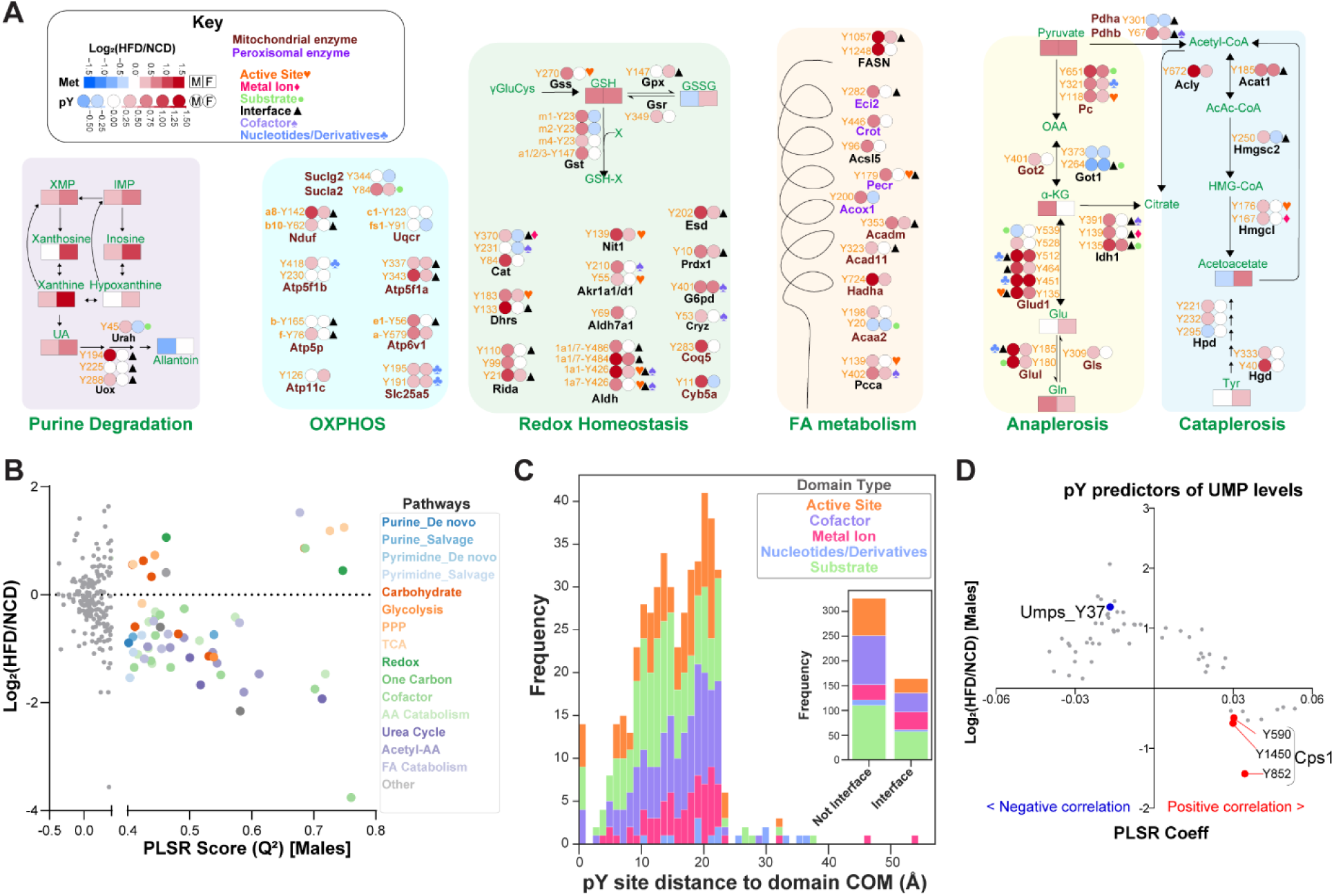
Integrative omics analysis yields pY sites that predict HFD-induced changes in metabolites. (A) Selected integrative map depicting pY sites and metabolites in pathways for purine degradation, oxidative phosphorylation (OXPHOS), redox homeostasis, fatty acid (FA) metabolism, and anaplerosis/cataplerosis. **(B)** Scatter plot of metabolite PLSR predictive score (Q^2^) vs fold change in Log2(HFD/NCD) where metabolites with valid model are color-coded by pathway. Each metabolite (Y-matrix) was regressed against the top 50% of pY sites with greatest change in magnitude (X-matrix) with k-fold cross-validation (see Methods). A model prediction score (Q^2^) ≥ 0.4 was used as an indicator of metabolites predicted by pY sites without overfitting. **(C)** Stacked histogram plot illustrating the frequency of predictive pY sites and their respective distances from functional domains and interface region. **(D)** Scatter plot of representative PLSR model coefficients vs Log2(HFD/NCD) of pY sites for uridine monophosphate (UMP), each datapoint is a pY site. PLSR coefficient indicates a positive (red) or negative (blue) relationship between UMP and the pY site. Highlighted sites are on enzymes in the *de novo* pyrimidine synthesis pathway: Umps Y37 and Cps1 Y590, Y1450, and Y852.

### Regression analysis identifies pY sites that predict changes in metabolites

In order to gain systems level insight and identify the covariation of metabolites with respect to pY on metabolic enzymes, we performed partial least squares regression (PLSR) analysis. Although PLSR was run for both sexes there were only a few metabolites predicted by pY for females due to the relatively low impact of HFD on these mice (Figure S4A). Diet induced changes were more pronounced in males, and therefore more metabolites were predicted by pY sites (Figure 4B, Table S8). Metabolites that met model predictive score (Q^2^) ≥ 0.4 included N-acetyl amino acids, cofactors, amino acid derivatives, nucleotides, and products of fatty acid degradation (Figure 4B). To identify the structural features of predictive pY sites, we selected sites with a variable importance in projection (VIP) score ≥1 and cross referenced them with structural annotations. Based on available structures, we annotated ∼65% of pY sites with respect to their domain proximity (Figure 4C, Table S9). Predictive pY were localized within 23Å of active sites, cofactor, and substrate binding domains (Figure 4C). A third of predictive pY sites were in both interface and functional domains of enzymes in amino acid metabolism, glycolysis/gluconeogenesis, TCA cycle, mitochondrial respiration, and fatty acid beta-oxidation (Figures 4C, S4B). For example, altered UMP levels are predicted by pY37 on Umps, which produces UMP, as well as multiple pY sites on the upstream enzyme carbamoyl phosphate synthase 1 (Cps1) (Figure 4D). PLSR model coefficient shows negative correlation of UMP with Umps pY37 and positive correlation of sites on Cps1 (pY590, pY1450, pY852), suggesting that pY37 could be an inhibitory site. Sites on Cps1 are in binding domains (Y1450) for its allosteric activator N-acetylglutamate (NAG) and ATP (Y590, Y852) ^75–77^. Thus, our PLSR model identified pY sites on known components of the *de novo* pyrimidine synthesis that are predictors of UMP levels. Overall, our integrative omics approach maps the systems level changes induced by HFD, identifies key pathways with sex specific pY response, and predicts the structural relevance of pY sites for enzyme function.

### Antioxidant supplements reverse HFD physiological phenotypes and molecular events

Systemic oxidative stress is prevalent in obesity^9,78,79^. Supplementing HFD with antioxidants abrogates the effect of obesity and metabolic syndrome^9,80–82^. Therefore, we asked whether the molecular events induced under HFD modeled by our PLSR analysis could similarly be reversed by supplementing HFD with exogenous antioxidant such as butylated hydroxyanisole (BHA). At the phenotypic level, BHA supplementation had a profound effect on HFD, especially in males. Compared to HFD alone, mice on BHA supplemented HFD weighed significantly less (FC ≈ 0.47) (Figure 5B), had a statistically significant decrease in fasted blood glucose levels (FC ≈ 0.47) (Figure 5C), and showed a ∼10-fold decrease in insulin levels (Figure 5D). To determine changes in metabolites and metabolic enzyme phosphorylation, we performed polar metabolomics and pY phosphoproteomics analysis on livers from mice on HFD with or without BHA supplement. We measured 258 metabolites across all samples, and found that BHA supplemented diet was anticorrelated with HFD (Figure 5E). PCA of the metabolites separates BHA supplemented samples from HFD only along the PC2 axis (Figure 5F). We then asked which metabolites were significantly altered in BHA supplemented HFD. The major drivers of separation on PC2 were in redox homeostasis, cofactor biosynthesis, amino acid catabolism, one carbon, and nucleotide metabolism pathways (Figure 5G). Specifically, metabolites in glutathione catabolism and recycling such as Ɣ-glutamyl-alanine, pyroglutamate/5-oxoproline, and ophthalmate were upregulated in BHA supplemented diets in both males and females (Figure 5G). Ophthalmate inhibits reactions that consume or efflux free GSH while promoting generation of cytoplasmic GSH, suggesting that BHA induced ophthalmate could be sustaining reduced cellular environment through management of the GSH pool^83^. End products of hexosamine biosynthesis pathways such as UDP-Glucose, UDP-GalNAc, UDP-xylose, and UDP-GlcNAc were also increased by BHA (Figure 5G). Moreover, utilization of urea cycle metabolites (citrulline to homocitrulline, arginine to argininic acid) was deceased by BHA, yet metabolites in the cytoplasmic arm of the urea cycle (argininosuccinate, citrulline, ornithine) and pyrimidines from *de novo* synthesis pathway were elevated. The drivers of differential clustering, therefore, indicate that BHA induces reprogramming of redox homeostasis, pyrimidine synthesis, and hexosamine byproducts while altering utilization of urea cycle metabolites.

**Figure 5.**
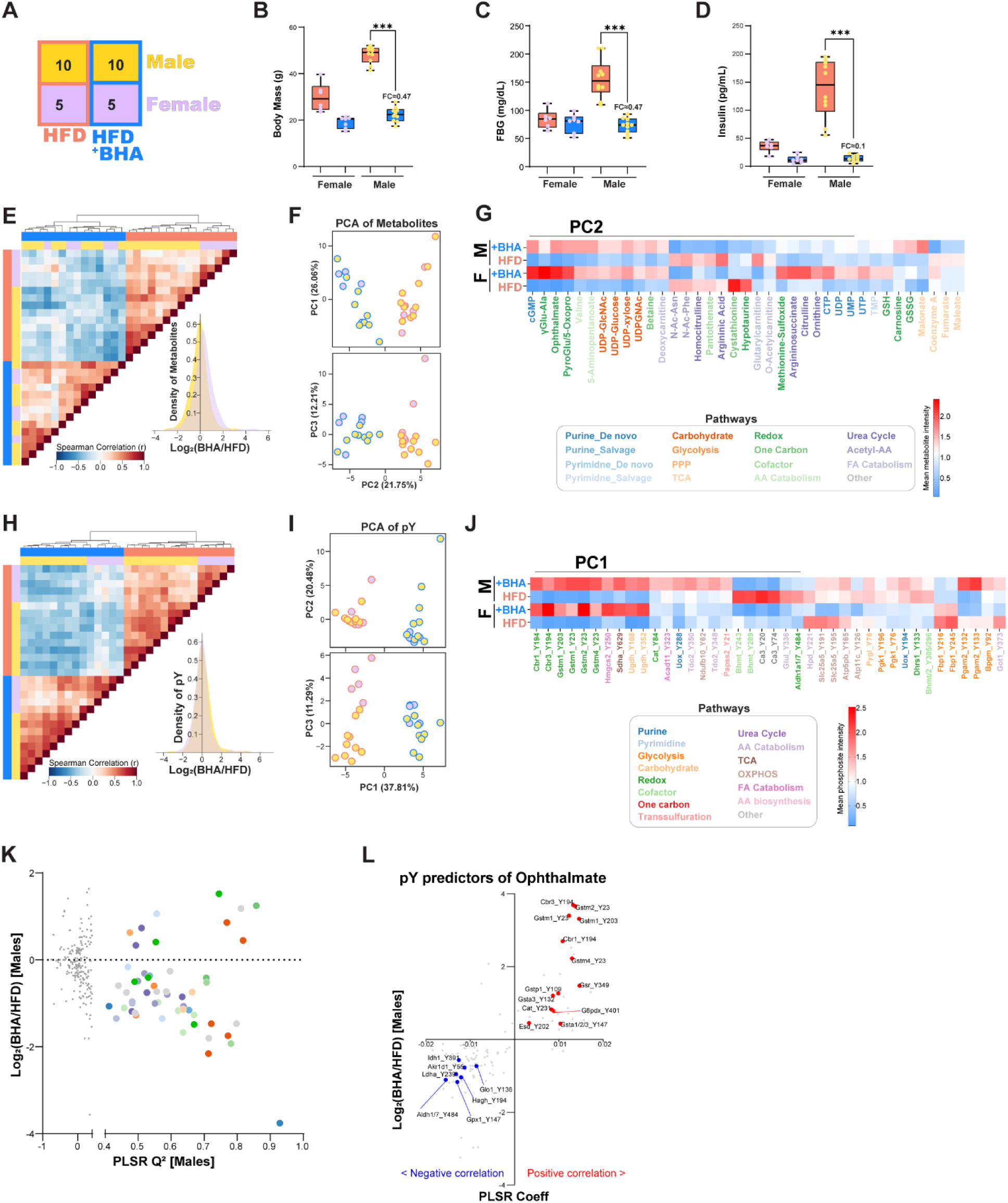
Supplementing HFD with the antioxidant BHA abrogates obesity phenotypes and rewires pY and metabolite signatures. (A-D) Experimental setup and physiological phenotypes of mice on HFD alone or HFD+BHA diet. Box and whisker plots with datapoints representing each animal. Two-way ANOVA with multiple comparison (Kruskal-Wallis test) was performed comparing corresponding HFD + BHA with HFD only groups and statistical significance was assigned: * p < 0.05, ** p < 0.01, *** p < 0.001. **(E)** Heatmap of spearman correlation analysis of polar metabolomics where columns and rows represent each animal. KDE of metabolite Log2(BHA/HFD) illustrating the variance in metabolite intensities between male and female samples. **(F)** Scatter plot of PCA for metabolite changes representing differential clustering along PC1, PC2, and PC3. Datapoints represent individual animals. **(G)** Heatmap of mean metabolite intensities altered by BHA and drive clustering along PC2, color-coded by pathway. **(H)** Heatmap of spearman correlation analysis and hierarchical clustering of samples based on pY phosphoproteomics. KDE of pY site Log2(BHA/HFD) representing variance between males and females. **(I)** Scatter plot of pY sites PCA where changes induced by BHA supplement are along PC1, PC2, and PC3. Data points represent individual animals. **(J)** Heatmap of mean phosphosite intensity of pY sites altered by BHA supplement and drive clustering along PC1. **(K)** Scatterplot of metabolites PLSR model score and fold changes, where metabolites with a valid score are color coded by pathway. PLSR analysis was performed with leave-one-out cross validation to identify pY sites (X-matrix) that predict metabolite (Y-matrix) response to BHA. **(L, M)** Scatter plot of pY sites for representative PLSR models for betaine and ophthalmate, where negative (blue) and positive (red) correlation of pY sites are indicated based on PLSR coefficient.

Targeted pY phosphoproteomics analysis yielded 231 pY sites on metabolic enzymes observed across all samples. Similar to metabolites, pY signatures were anticorrelated between HFD alone and BHA supplemented HFD, but the difference between males and females was not significant (Figures, 5H, 5I). BHA altered pY on numerous enzymes with most of them belonging to redox homeostasis, cofactor biosynthesis, one carbon metabolism, oxidative phosphorylation (OXPHOS), and fatty acid metabolism (Figure 5J). For example, pY on NADH/NADPH generating enzymes such as Ldha, Idh1, G6pdx, Aldh1a1/7, Akr1a1/d1, Cat, Esd, and Gsr were localized in close proximity to cofactor binding domains and were reversed by BHA supplement (Figure S5A). Additionally, pY on glutathione binding enzymes such as Gst, Cbr, Gpx1, and Gsr were further increased by BHA (Figure S5B). Overall BHA-induced pY changes in HFD were concentrated on redox enzymes that cycle NADH, NADPH, and GSH.

PLSR analysis identified BHA-modulated predictive pY sites were in urea cycle, redox homeostasis, and glycolysis (Figure 5K, Table S10). For example, BHA-induced changes in ophthalmate are positively correlated with pY sites mostly on GSH consuming enzymes (e.g., Gstm/p, Cbr1/3), and negatively correlated with pY sites on NADH/NADPH cycling enzymes (e.g., Idh1, Akr1d1, Ldha, Gpx1) (Figure 5L). Since ophthalmate has been shown to regulate GSH consumption, it is possible that sites on GSH consuming enzymes could be inhibitory, while sites on NADH/NADPH cycling enzymes could be activating. Together, these results demonstrate that supplementing HFD with small molecule antioxidants can ameliorate obesity while altering the molecular events associated with redox homeostasis and metabolic syndrome.

### CRISPRi-based validation of pY sites and their role on enzyme activity

Our PLSR computational models highlighted associations between metabolic enzyme phosphorylation sites and altered metabolites in HFD and HFD+BHA conditions while also predicting the functional role of multiple sites found in specific structural domains. To validate these predictions biochemically, we developed an approach using doxycycline (Dox)-inducible CRISPRi^84,85^ with overexpression rescue of enzyme variants to assay activity in A549, a human lung cancer cell line that strongly expresses redox enzymes. Based on our multiomics data and predictive computational models, as well as previously published work, we selected GSTP1 as a candidate enzyme to validate our CRISPRi-rescue approach^86–89^ (Figure 6A). GSTP1 phosphorylation on multiple pY sites is significantly decreased in HFD compared to NCD, especially in males, and BHA supplements reversed HFD induced loss of GSTP1 pY. In many of our PLSR models, using both targeted and untargeted phosphoproteomics data, we were able to capture pY sites on GSTP1 as predictors of various metabolites (e.g., argininic acid, pyroglutamate/oxoproline, betaine, ophthalmate) (Figure S6A). Moreover, GSTP1 is highly conserved across mammals, and the phosphorylation sites we measured are conserved in various GST isoforms^89–91^. To determine the functional consequence of pY sites on GSTP1, we first knocked down GSTP1 (Figure S6B), then stably expressed Dox-inducible GSTP1 WT, phosphodeficient (Y→F) or phosphomimic (Y→E) mutants for both well-established and understudied pY sites: Y8, Y64, Y109, Y199 (Figure 6A). Using 1-chloro-2,4-dinitrobenzene (CDNB) decay as a readout, we performed cell-based GSTP1 activity assays. Compared to wild type (WT), Y8F and Y8E both showed reduced activity. Y8 is a highly conserved and well-established active site residue, and loss of this active site in any capacity renders the enzyme inactive (Figures 6B, S6C, Table S11). Loss of phosphorylation at Y64 and Y109 via Y→F mutations reduced activity by ∼20-30% compared to WT, but there was still substantial activity compared to GSTP1 knockdown (mCh). However, Y64E and Y109E both had significantly (p<0.001) reduced activity comparable to GSTP1 knockdown (Figure 6B, Table S7). Lastly, Y199F and Y199E both had lower activity compared to WT by ∼30%, but not as low as GSTP1 knockdown. Therefore, our cell-based assays suggest that pY at sites Y64 and Y109 are inhibitory, while phosphorylation of Y199 show minimal impact on GSTP1 activity. To further validate our CRISPRi-rescue experiments, we designed an *in vitro* assay using recombinant GSTP1 variants coupled to hyperphosphorylation with recombinant SRC kinase (Figure 6C). In agreement with the results from our CRISPRi-rescue experiment, Y8F, Y8E, Y64E, and Y109E all showed significant decrease in activity compared to WT in both +/- SRC conditions (Figures 6D, S6E, Tables S12, S13). SRC kinase phosphorylated GSTP1 variants overall showed modest decrease in activity with Y199E showing statistically significant decrease compared to the corresponding no SRC condition. Therefore, our CRISPRi-rescue approach was able to capture enzyme activity of GSTP1 variants in a similar manner to our orthogonal *in vitro* approach, thus cross-validating our observations. Overall, these results show that phosphorylation of GSTP1 on Y64 and Y109 are inhibitory, and as previously published, Y8 is an essential active site tyrosine that is indispensable for GSTP1 function regardless of gain or loss of phosphorylation status^89^. We also noted that the phosphomimic mutant Y199E showed significant decrease in activity when incubated with SRC, suggesting that hyperphosphorylation might be a negative regulator for GSTP1 as a whole.

**Figure 6.**
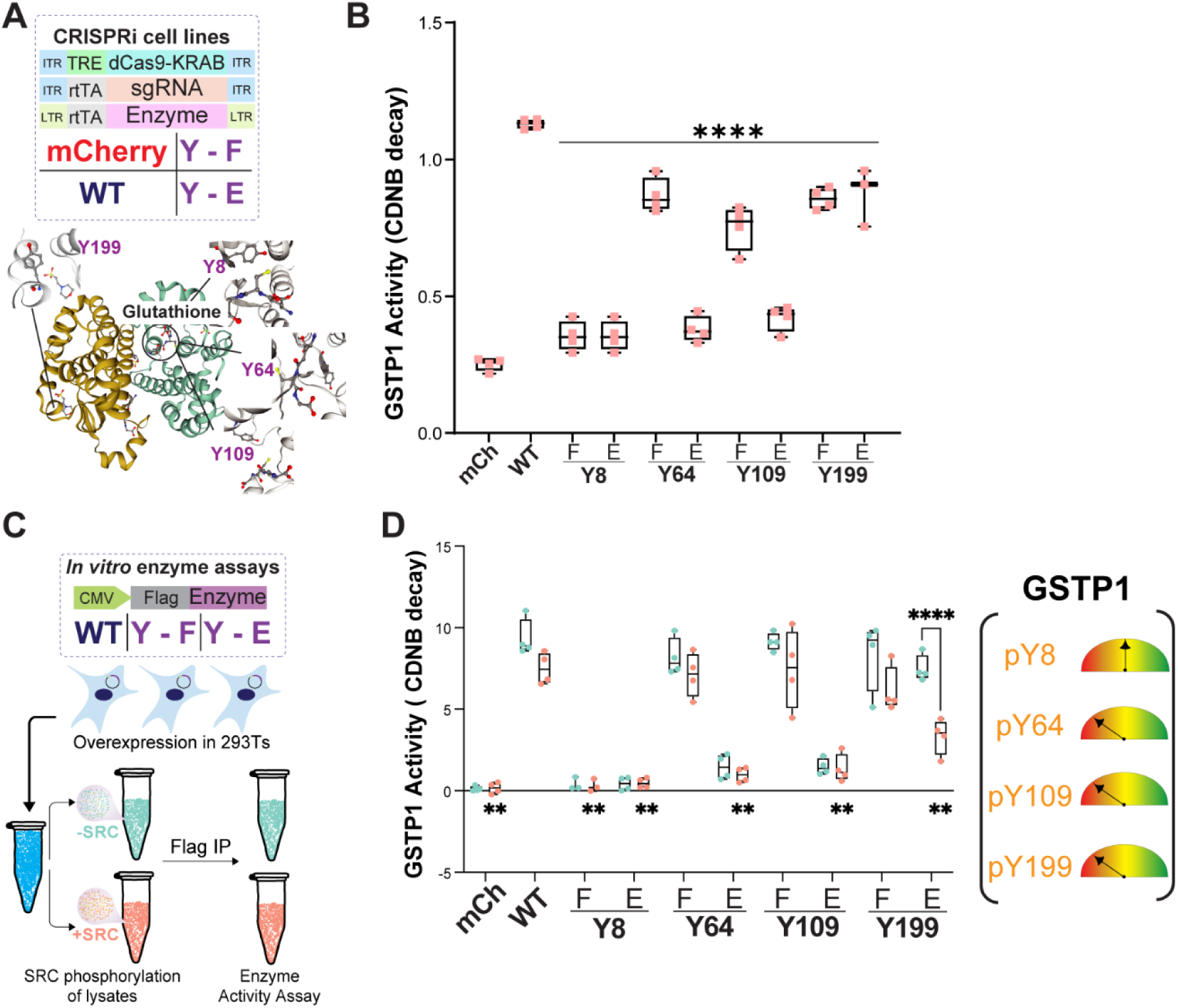
CRISPRi-rescue validates pY sites on GSTP1 as inhibitors of enzyme activity. **(A)** Schematic of Dox inducible CRISPRi-rescue for cell-based assays and select pY sites on GSTP1 selected for benchmark (PDB: 5gss). **(B)** Box and whisker plot representing CDNB day assay on A549 cells treated with doxycycline for 96 hours to measure activity of GSTP1 variants (N=4). **(C)** Schema of orthogonal approach where recombinant Flag tagged GSTP1 variants were overexpressed in HEK293T cells, and the lysate was divided into control or recombinant SRC kinase-hyperphosphorylated samples before Flag immunoprecipitation and enzyme activity assay. **(D)** CDNB decay assay was performed on 1µg of recombinant protein purified from SRC-hyperphosphorylated lysates. mCh=mCherry, WT=wildtype. Statistical analysis was performed using two-way ANOVA with multiple comparison (Dunnett test) of phospho-variants to wild type (WT). For in vitro assays comparisons were done across all +SRC conditions, as well as -SRC and +SRC conditions within each variant. Statistical significance was assigned as follows: * p < 0.05, ** p < 0.01, **** p < 0.0001.

### CRISPRi-rescue coupled with isotope tracing metabolomics confirms IDH1 Y391, UMPS Y37 as activity regulating phosphosites

As further experimental validation of our computational model predictions, we coupled our inducible CRISPRi-rescue system with stable isotope-tracing metabolomics. We focused on pY sites with differential response to HFD, those with potential structural role gathered from annotation, sites modulated by BHA, and predictive sites from our PLSR models. We selected both previously characterized (IDH1 Y391) and understudied (UMPS Y37) sites of interest identified in our pY analysis. We then asked whether these phosphorylation sites affect enzyme function and pathway kinetics. Since the pathways/enzymes of interest were downstream of glutamine anaplerosis, we performed U-^13^C5 Glutamine stable isotope tracing in IDH1 or UMPS Dox-inducible CRISPRi-rescue of the appropriate WT, Y→F, or Y→E variants.

IDH1 Y391 is localized in the interface of two-monomers and NADP binding domain ^92^ (Figure 7A). Moreover, pY391 is upregulated by HFD and downregulated by BHA in males where it emerges in many of our PLSR models as a negatively correlated predictor of redox metabolites (e.g., pyroglutamate/oxoproline, ophthalmate). Phosphorylation at Y391 has been established as an activator phosphosite that promotes reductive carboxylation of alpha-ketoglutarate (aKG) into citrate in the cytoplasm, especially in the context of defective mitochondria^93–101^. Therefore, we performed kinetic glutamine tracing for 1h and 2h then measured citrate generated via oxidative pathway in the TCA cycle (citrate M+4) or reductive pathway from aKG (citrate M+5) (Figure 7A). Knockdown of IDH1 increased citrate M+4 and reduced citrate M+5, suggesting that decreasing IDH1 levels promotes more oxidative pathway via TCA cycle (Figure 7B, S6A, Table S14). Both WT and Y391F had comparable M+4 and M+5 citrate, but Y391E decreased M+4 citrate while increasing M+5 citrate. Moreover, M+3 aspartate generated from citrate in the reductive pathway was also elevated in Y391E expressing cells. However, the total pool size, or the sum of labeled and unlabeled species, of neither citrate nor aspartate were significantly altered across all conditions (Figure 7C). These data confirm that phosphorylation at Y391 induces IDH1 activity promoting reductive carboxylation (Figure 7D), thereby increasing levels of reductive carboxylation-derived downstream metabolites such as aspartate that feed into nucleotide synthesis. Moreover, downregulation of pY391 by BHA in males suggests that there is engagement of reductive metabolism dependent on the severity of obesity phenotypes and metabolic syndrome.

**Figure 7.**
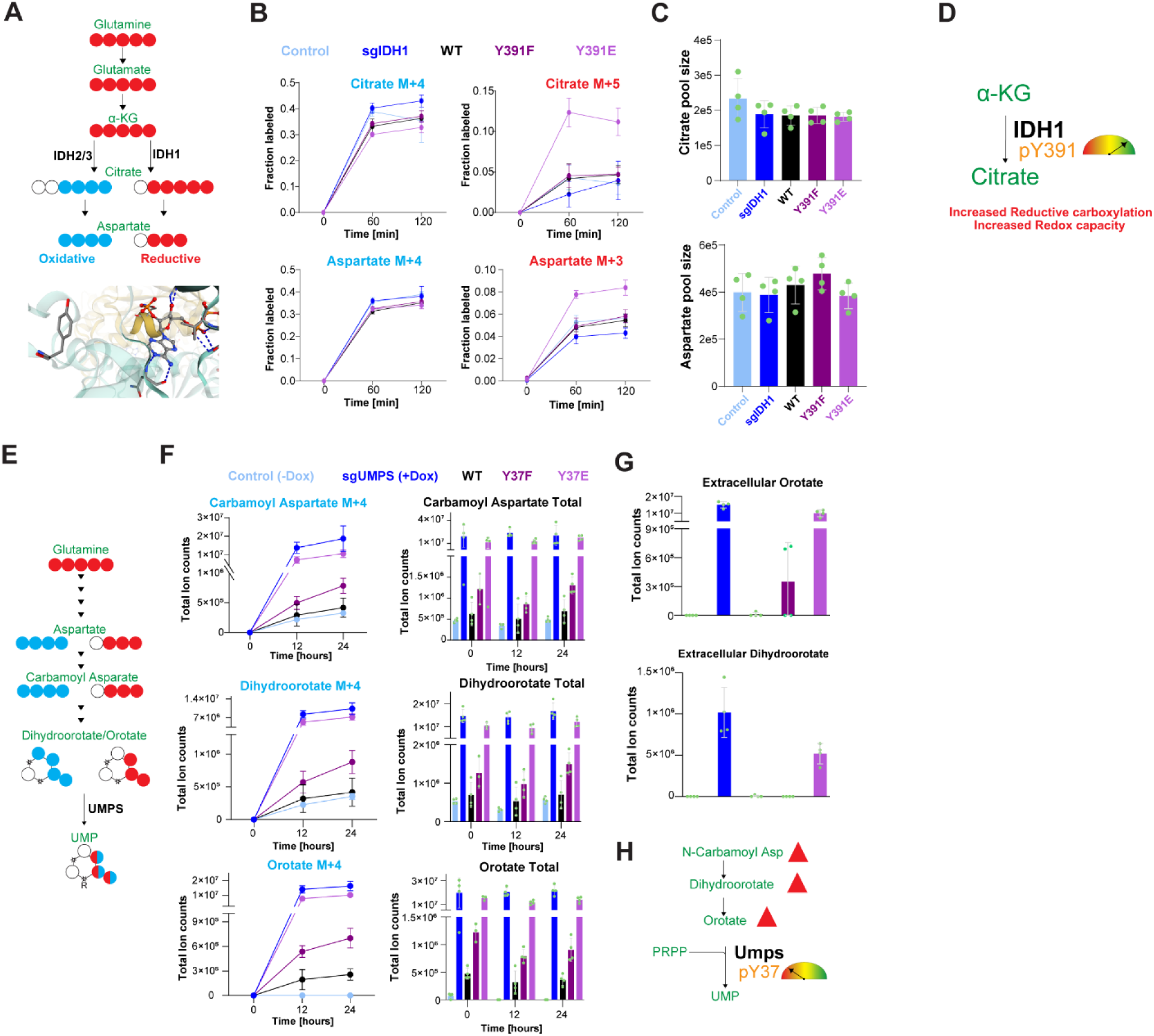
Interrogation of pY functional role by coupling isotope labeled tracing and CRISPRi-rescue validates regulatory role of IDH1 Y391 and UMPS Y37 on enzyme activity. (A) U-^13^C5-Glutamine stable isotope tracing scheme to evaluate the role of IDH1 Y391 which is positioned in the NADP binding domain (PDB:1t0l)^92^. **(B)** Fraction labeling of citrate and aspartate via oxidative (blue) and reductive (red) TCA cycle over 0, 60, or 120min in A549 sgIDH1 CRISPRi-rescue cell lines (N=4). Knockdown-rescue was achieved by adding 500ng/mL of Dox, replenished daily, to culture media for a total of 96 hours. Total ion counts were normalized to Norvaline internal standard and total protein content, and fraction labeling was calculated by taking the ratio of each labeled species divided by the total pool size. **(C)** Pool sizes of citrate and aspartate at 120min (N=4). Pool sizes were calculated by summing all species of each metabolite per condition. Data available in Table S8(A – C). **(D)** Model showing pY391 activating IDH1 to induce reductive carboxylation of aKG to citrate. **(E)** U-^13^C5-Glutamine stable isotope tracing scheme for 0, 12, or 24 hours to measure changes in *de novo* pyrimidine synthesis pathway in A549 sgUMPS CRISPRi-rescue cell lines treated with dox for 96 hours total. **(F)** Total ion counts of M+4 labeled and total *de novo* pyrimidine synthesis intermediates carbamoyl aspartate, dihydroorotate, and orotate (N=4). **(G)** Extracellular orotate and dihydroorotate were measured from media collected at the 24 hours timepoint. Data available in Table S8(D – G). **(H)** model for pY37-mediated inhibition of UMPS enzyme activity.

UMPS is a bifunctional enzyme that carries out the final step in *de novo* pyrimidine biosynthesis where UMP is generated in a two-step reaction^59–61^. UMPS Y37 is found in the N-terminal portion of the enzyme (OPRTase) that transfers phosphoribosyl group onto orotate to create OMP, which is subsequently converted to UMP by the terminal reaction performed by OMP decarboxylase (ODC) located in the c-terminal domain^102^. Phosphorylation at Y37 has been identified many times in the published phosphoproteome, but there has not been any characterization of how this site affects *de novo* pyrimidine synthesis and peripheral pathways. Structural annotation shows that Y37 directly interacts with OMP via pi-stacking and the hydroxyl group forms a water bridge with the orotate ring (Figure 2D)^102–105^. UMPS Y37 is highly phosphorylated in males on HFD, and our PLSR model shows negative correlation between UMP and pY37, suggesting that this site might serve an inhibitory role in *de novo* pyrimidine synthesis. Moreover, pY37 is reduced by BHA supplementation, and *de novo* pyrimidine synthesis pathway metabolites are upregulated by BHA, especially in females (Figure 5G). To validate our *in vivo* observation for UMPS pY37, we traced U-^13^C5 Glutamine incorporation into *de novo* pyrimidine synthesis intermediates for 12h and 24h (Figure 7E). Glutamine carbons are incorporated into aspartate which then enters *de novo* pyrimidine synthesis as carbamoyl aspartate^60^. Subsequently carbamoyl aspartate is converted into dihydroorotate then orotate, the starting substrate used by the OPRTase^59^. Therefore, glutamine labeled metabolites upstream of UMPS (carbamoyl aspartate, dihydroorotate, and orotate) should have either 3 (M+3) or 4 (M+4) labeled carbons (Figure 7E). Compared to control, Dox-induced knockdown of UMPS leads to build up of carbamoyl aspartate, dihydroorotate, and orotate (Figures 7F, S7B, S7C, Table S14). Both labeled and total pools of metabolites upstream of UMPS were significantly accumulated in UMPS knockdown and Y37E expressing cells, while cells overexpressing Y37F and WT had similar phenotype to control. The phosphoribosyl conjugated orotate, orotidine, was only elevated in knockdown cells but not in cells expressing UMPS variants (Figure S7D). These data suggest that the negative charge introduced by phosphorylation of UMPS Y37 and not the aromaticity of the residue leads to decreased enzyme activity. Moreover, analysis of the media collected after 24 hours shows that the backed-up *de novo* pyrimidine intermediates efflux into the extracellular space (Figure 7G). We do not observe an appreciable difference in UMP, UDP, UTP, and CTP levels (Figure S7E), however this is not surprising since recycling pathways can sufficiently replenish the nucleotide pool. In agreement with our PLSR predictions, these results show strong evidence that phosphorylation of UMPS at Y37 leads to decreased enzyme activity. This phenotype is akin to defects in *de novo* pyrimidine synthesis observed in orotic aciduria as well as excess uric acid which inhibit the ODC domain of UMPS^106,106–108^. Thus, we can infer that pY37 could be preventing transfer of orotidine to ODC for the final step of UMP synthesis thereby creating a bottleneck in *de novo* pyrimidine synthesis. Moreover, it has been established that high orotate levels induce fatty liver/steatosis via *de novo* lipogenesis in model organisms and human cell lines^109–111^, suggesting a mechanism on how UMPS pY37 and other phosphosites contribute to HFD-induced phenotype as part of the concerted network of metabolic enzyme phosphoproteome in obesity.

## Discussion

In this study we used a structural and multiomic approach to annotate, evaluate, and characterize how phosphorylation of metabolic enzymes, specifically pY, alter metabolic activity. One of the challenges of characterizing phosphosites on metabolic enzymes is the lack of systems level data that can elucidate which phosphosites are critical for enzyme function specifically following perturbations. Here we integrated public domain knowledge of enzyme structures to identify phosphosites with potential roles in enzyme function. In an effort to structurally contextualize phosphosites, we measured their proximity to functional domains. Our results illustrate that the metabolic phosphoproteome is enriched for oxidoreductases with phosphosites mainly localizing in binding domains for substrate, cofactor, and active sites. Despite low cellular stoichiometry, pY were the most enriched phosphosites in functional domains. A significant proportion of pY in active sites serve as transition state stabilizers or proton donors (e.g., IDH1 Y135/139, AKR1C1/A1 Y55, PGAM1 Y92), which underscores that pY sites present unique structural features and serve as signaling input into direct modulation of enzyme function. In an effort to comprehensively evaluate the structural features of phosphosites, we also used the PISA analysis tool to identify published phosphosites localized in dimerization interfaces. Again, pY were the most prominent phosphosites in dimerization domains that play a significant role in complex formation. Dimerization interactions between phosphosites and surrounding residues favor the utilization of the hydroxyl group on the phosphosite, especially for pY; thus, addition of a phosphorylation moiety impacts enzyme dimerization. For example, LDHA Y10 is localized in the interface domain, and phosphorylation promotes formation of an active tetrameric enzyme^33,36,112^. From a number of published LDHA structures we were able to identify that the OH group on LDHA Y10 makes contact with either glycine, glutamate, or lysine residues while the backbone makes contact with leucine. We find that dimerization interface phosphosites preferentially interact with charged or polar residues, which could be the key determinant for what role these sites serve when phosphorylated. The potential impact of phosphorylation in interface regions could be increased hydrophobicity, destabilizing/stabilizing interactions, and altered conformation that can all lead to altered enzyme function. Future work could interrogate what common features or interactions define residues that promote versus deter dimerization.

The role of phosphosites on metabolic enzymes are context dependent as different perturbations result in varying signaling and metabolic signatures. Using multiomics and computational modeling, we illustrate that HFD induced, sex-specific physiological changes are associated with phosphoproteomic and metabolomic alterations. The majority of the differences between sex and diet conditions were in redox homeostasis, fatty acid, nucleotide, and amino acid metabolism. For example, females exhibit increased phosphorylation of enzymes involved in waste elimination pathways such as ammonia and urea. It is possible that the severity of HFD phenotypes could be dictated by engagement of waste elimination processes that maintain mitochondrial health. Phosphosites on enzymes involved in reductive metabolism (e.g., IDH1 Y391) are upregulated in males on HFD, as it has been established, reductive metabolism is engaged during hypoxia, oxidative stress, and mitochondrial dysfunction^93,95,97,98,113^. This is especially true during *de novo* lipogenesis and glutamine/glucose stimulated insulin secretion^98,114–116^. Our *in vivo* model demonstrates that the sex specific differences we observe in obese conditions are partly driven by adaptive response that engage reductive metabolism. Accordingly, treating mice on HFD with antioxidant (BHA) abrogated the physiological effects of HFD and altered both redox metabolites as well as pY site on antioxidant enzymes. Additionally, changes incurred by BHA supplement underscore that sex difference in obesity stem from capacity to store fats and to detoxify ROS/RCS/RNS. For example, phosphorylation on GSH or NADH/NADPH cycling enzymes show an adaptive response to BHA, and thus might be associated with increased fatty acid oxidation which is known to induce oxidative stress^21,117,118^. As a result, altered amino acid metabolism could be a mechanism to replenish oxidative reactions in the TCA cycle via anaplerosis (e.g., pyruvate, glutamine).

HFD results in hyperphosphorylation of enzymes involved in glutamine anaplerosis as well as antioxidant enzymes that detoxify RCS (Cbr1/3, Aldh1a1/7, Aldh2 etc.) to a greater degree in male mice. Increased phosphorylation of metabolic enzymes in males, where there is severe phenotype, suggests that most pY sites induced by HFD could serve as constraints to rein in defects in redox balance. Our data provides evidence that more cytoplasmic redox reactions are activated in BHA conditions (e.g., elevated ophthalmate, GSSG). Additionally, phosphorylation of methionine and bicarbonate producing enzymes (Bhmt, Ca3) is downregulated by BHA further supporting the redox homeostasis is at the center of HFD induced changes that might be responsible for obesity phenotypes^119–122^. Antioxidation via BHA supplement reverses most of the HFD induced pY on enzymes. However, this is a double-edged sword as chronic antioxidation can also result in reductive stress^123–127^. High dose antioxidant supplements such as NAC and Vitamin E cause accumulation of reductive stress in brown adipose tissues by increasing mitochondrial ROS that results in dysfunction^127^. The effect size on HFD in our study could be because the antioxidation mechanism for BHA (donates hydrogen to oxidized molecules) is distinct from NAC (alters cellular cysteine pools) and Vitamin E (reduces lipid peroxidation). Additionally, mitochondrial specific antioxidants such as CoQ10 have been effective at directly targeting hyperglycemia induced redox imbalance and mitochondrial dysfunction^82,117^. Therefore, future work should explore the sex specific effects of different antioxidants in obesity, and strategies to strike redox balance without skewing towards reductive stress. We can circumvent complications resulting from excessive antioxidation by monitoring the currencies of redox homeostasis such as NADH/NADPH, GSH, TRX, Cysteine, FMN/FAD, and acetaldehyde/alcohol which provide high reducing potential and could indicate sexually dimorphic redox imbalance^128,129^. Coordination of insulin secretion by metabolic coupling factors (MCFs) such as ATP, NADH, glucose, glutamate, long chain acyl coenzyme A, and diacylglycerol can also serve as an indicator of systemic dysregulation^130^. In combination with measurement of pY sites on enzymes that cycle the aforementioned reducers, we can construct a comprehensive network level view of redox homeostasis in the development of obesity and progression towards metabolic syndrome.

As we work towards overcoming technical challenges of detecting pY on metabolic enzymes with more targeted approaches, as done here, we need to carefully design extraction protocols for metabolite and protein. There are a number of new approaches to extract metabolites and proteins within the same workflow. While a combined extraction method would be useful for comparisons across different layers of biomolecules, such extraction methods may affect how well pY are preserved throughout the sample processing stage. Due to the low abundance and rapid dynamic regulation of pY^3^, we opted to use parallel extractions by dividing the sample between metabolomics and phosphoproteomics. We believe our approach prioritizes sample preservation while maintaining rigor and reproducibility via quantitative omics.

Our CRISPRi-rescue approach which shifts the stoichiometry of endogenous enzyme to phospho-variants, elucidated a number of functionally relevant pY sites on GSTP1, IDH1, and UMPS. We used previously characterized pY sites as benchmark to validate our method and identified that GSTP1 Y64 and Y109 inhibit enzyme activity. We also note that hyperphosphorylation of GSTP1 could serve as an overall inhibitory control system. In contrast, phosphorylation of Y391 on IDH1 validated as an activator of enzymatic function and promoter of reductive carboxylation. Reductive carboxylation allows cells with defective mitochondria to maintain redox balance^94,97,98,113^. If HFD induces mitochondrial stress due to increased oxidative reactions, IDH1 pY391 and other sites might facilitate redox metabolism reprograming to maintain homeostasis during increased fatty acid oxidation. Therefore, reductive metabolism is essential for redox balance in chronic obese conditions. Our analysis of UMPS establishes the inhibitory role of pY37, which could be serving as the gatekeeper between the bifunctional enzyme domains: OPRTase and ODC. We hypothesize that pY37 creates a large, negatively charged region to prevent OMP cycling, reducing enzyme efficiency, thus creating a bottleneck that causes the buildup of upstream metabolites carbamoyl aspartate, dihydroorotate, and orotate. This aligns with observations that inhibition of ODC by excess uric acid or other inhibitors leads to a buildup of carbamoyl aspartate, dihydroorotate, and orotate^102–105^. Furthermore, increased *de novo* pyrimidine synthesis metabolites can create an undue burden on the mitochondria because ROS production and quinone reduction is strongly tied to oxidation of dihydroorotate to orotate^106,106–108^ ^109–111^. Thus it is not surprising that, similar to our BHA study, treating HFD induced obesity in mice with carnitine orotate complex (Godex) or carnosine analogs have been shown to effectively mitigate obesity, inflammation, and insulin resistance ^81,111,131–133^. We also observed that carnosine and anserine are decreased in males compared to females treated with HFD, thus underscoring that sex-differences in obesity might arise from the potential to detoxify reactive molecules generated by mitochondrial oxidative metabolism. Future studies could explore how these differences in detoxification potential alter electron transport, mitochondrial levels of reducing equivalents, and how combined pY on multiple enzymes contributes to the OSR metabolon. An interesting observation from our biochemical interrogation of pY sites in the aforementioned enzymes is that the regulatory role of phosphosites was specifically linked to the negative charge and not the aromaticity of the residue since Y→F mutations were similar to WT. Therefore, exploring when and how the charge state versus the aromaticity of phosphosites alters activity can provide valuable insights into the key structural features of pY sites on metabolic enzymes. While our CRISPRi-rescue approach does not eliminate the endogenous enzyme completely, it allows for a well-controlled stoichiometric shift to the variant of interest. Our observations for GSTP1, IDH1, and UMPS all underscore that shifting the stoichiometry of modified enzymes can have significant ramification for the corresponding pathway and for peripheral pathways that can amplify the change into systemic metabolic tuning. We believe this provides a more realistic way to examine the impact of PTMs on metabolic enzymes compared to, for example, ablating endogenous variants via knockout. Subtle, yet concerted additive coordination of PTMs on multiple components of a given metabolic pathway can create directed dynamic changes to reach a more thermodynamically favorable state for sustained chronic metabolic reprogramming.

## Study limitations

Our comprehensive computational interrogation of the structural features of phosphosites provides a unique approach, but we are limited by the currently existing knowledge base for both structural data and the published phosphoproteome. Phosphorylated metabolic enzymes are not very well represented on published structures: thus, our analysis has focused on the structures of the apo enzymes to identify potential function. Conformational changes due to protein phosphorylation are missing from our analysis because phosphorylated structures have not been determined. Molecular dynamics simulation analysis of sites on a few enzymes (e.g., PGAM1, G6PD)^134,135^ illustrate how the dynamics and structure of metabolic enzymes are altered upon phosphorylation; however, this is a computationally demanding endeavor to apply on a large scale. As technology improves, we anticipate a wider application of this strategy to characterize the structure-function relationship of phosphosites on metabolic enzymes. Indeed, the advent of AlphaFold and complex deep learning models will likely increase our capacity to glean biochemical insight^136^. Similarly, we are limited with the currently available technology for mass spectrometry-based proteomics and metabolomics, which are rapidly evolving^137^. Due to the low abundance of pY sites on metabolic enzymes, they are often masked by higher abundance sites on signaling, RNA binding, and cytoskeletal proteins, among others.

Quantitative targeted approaches, as done here, can be a viable solution to assess how perturbations alter phosphorylation on metabolic enzymes. Additionally, metabolite identifications can be optimized by increasing sensitivity and incorporating labeled standards to better resolve biologically relevant small molecules and improve input data for computational modeling. By overcoming these limitations, we will be able to see deeper and clearer into the complex network of signaling-mediated metabolic regulation.

## Methods

### Structural annotation of phosphosites

Published phosphorylation sites were retrieved from the PhosphoSitePlus (PSP) database (version Dec 2023)^40^. Metabolic enzyme dataset was obtained from the Kyoto Encyclopedia of Genes and Genomes (KEGG)^41,42^. Gene level pathway enrichment analysis was performed using Enricher API and Reactome 2022 and Go Molecular Function 2023 gene sets^43–45^. In order to annotate phosphosites on metabolic enzymes, published phosphosites were filtered to only include proteins with identifiers for enzyme classification (EC), KEGG, and those that have published structures in the Protein Data Bank (PDB)^46^. All available structures were obtained from RCSB PDB, and structural quality determined based on resolution (≤2.6Å ) and Rfree – Rwork (≤0.05)^47^. X-ray structures meeting the defined cutoff were then used for further phosphosite characterization. For each protein with high quality structure(s), we curated binding domain information from features annotated on Uniprot-KB^48^. We then used PyMol (The PyMOL Molecular Graphics System, Version 3.0 Schrödinger, LLC) to calculate the center-of-mass (COM) for each domain on our list of proteins, and measured the distance between any known phosphosites and the domain COM. Hypergeometric distribution test was performed as follows: sample success = phosphosites within 23 Å of domain, sample size = all unique serine (S), threonine (T), or tyrosine (Y) residues within 23 Å, population success = all phosphosites on proteins with domain of interest, population = all unique S, T, or Y in proteins with domain of interest. Multiple hypothesis correction of p-value from the cumulative distribution function was performed using Benjamini-Hochberg method (α=0.05). Dimerization domains were separately annotated using PDBePISA^49–51^. Using the PISA API we retrieved the interface information for all multimeric structures, and both residue interaction as well as property information was extracted to generate the interface dataset. Example structures and images were generated using SwissModel^103,104,138^, PLIP^105^ tool, and the following PDBs: AKR1C1 (1mrq)^139^, ACAT1 (2iby)^140^, G6PD (7sni)^55^, IDH1 (1t0l)^92^, UMPS (2wns)^102^. All data used for structural annotation can be accessed in Table S1.

### Mouse work

C57BL/6J mice were obtained from The Jackson Laboratories and housed in a specific pathogen-free facility accredited by the Association for Assessment and Accreditation of Laboratory Animal Care using laminar flow cages at 21°C. Mouse health was monitored and maintained by veterinary staff under the supervision of a veterinarian. Male and female mice fed *ad libitum* either a standard chow diet (NCD, F4031, Bio-Serv), a high fat diet (HFD, S3282, Bio-Serv), or a HFD supplemented with 1.5% 2(3)-tert-butyl-4 hydroxyanisole (BHA, S7958, Bio-Serv) for 16 weeks beginning at age 8 weeks. All mice were euthanized at age 24 weeks following an overnight fast by inhalation of isoflurane followed by cervical dislocation, and the livers were snap frozen in liquid nitrogen upon removal. All experiments were carried out in accordance with guidelines for the use of laboratory animals and were approved by the Institutional Animal Care and Use Committees (IACUC) of the University of Massachusetts Chan Medical School.

### Protein digestion, Cleanup, and Multiplexing

Pulverized mouse livers were homogenized in 8M urea and lysates were cleared by centrifugation at 21,000xg at 4^°^C. Protein concentration of the cleared lysates was determined using bicinchoninic acid assay (BCA assay). 1mg of protein was reduced by adding 10mM DTT at 56^°^C for 1 hour and alkylated with 55mM IAA at room temperature protected from light for 1 hour. Samples were diluted 10-fold (v:v) with 100mM ammonium acetate (pH=8.9) and digested using sequencing grade Trypsin (1:50 w:w, 20µg Trypsin : 1mg protein), rotated overnight at room temperature. Digestion was quenched with 1:20 (v:v) of 100% glacial acetic acid. Peptide desalting and cleanup was performed using SepPak Light C-18 cartridges (Waters, WAT023501) attached to 10mL syringes on a syringe pump. Briefly, cartridges were washed at 2mL/min using 0.01% acetic acid, then 90% acetonitrile in 0.01% acetic acid, and equilibrated with 0.01% acetic acid before loading peptides at 0.8mL/min. Peptide bound cartridges were washed with 0.01% acetic acid, and peptides were eluted in 40% acetonitrile with 0.01% acetic acid. The elution was then concentrated via speed vac and peptide BCA assay was performed to determine concentration. For normalization across multiple runs, an independent sample of linear combination of all samples (i.e.,, a bridge) was generated. Then 200µg peptides per sample were lyophilized for multiplexing. Dried peptides were resuspended in 50mM HEPES (pH=8.5) and multiplexed using TMT 16plex reagent at 200µg:490µg (peptide:TMT). Labeling was carried out for 4 hours at room temperature then quenched with 1:15 (v:v) of 5% hydroxylamine for 15min before combining all samples. To ensure maximum peptide recovery and reaction quenching, sample tubes were washed twice with 25% acetonitrile in 0.01% acetic acid and the washes were also collected to the sample mixture. Labeled peptides were dried down via speed vac, and stored at -80^°^C before pY immunoprecipitation.

### Immunoprecipitation, phosphopeptide enrichment, and liquid chromatography

Dried TMT labeled peptides were resuspended in IP buffer (100mM Tris-HCl, 1% Nonidet P-40, pH=7.4), and pY peptides were immunoprecipitated overnight using Protein G agarose beads conjugated to 24µg of Super-4G10^141,142^ and 6µg of PT-66 anti-pY antibodies. Supernatant of pY IP was stored for crude protein analysis, and beads were washed with IP buffer. Peptides were eluted twice from beads using 0.2% trifluoroacetic acid at room temperature, and the elution was further enriched using High-Select Fe-NTA phosphopeptide enrichment spin columns. Enriched phosphopeptides were eluted into a BSA coated 1.7mL tube, and dried down via speed vac. Enriched phosphopeptides were then resuspended in 5% acetonitrile with 0.1% formic acid solution and directly loaded on to an analytical column (inner diameter= 50µm) packed with 3µm C18 beads. Peptides were separated via liquid chromatography (LC) with 150 minutes gradient of 0 to 70% acetonitrile with 0.01% acetic acid (Buffer B) and 0.01% acetic acid (Buffer A) at a flowrate of 0.2mL/minute on an Agilent 1260-Infinity LC coupled to Ortibitrap Exploris 480 Mass Spectrometer. Chromatography settings for % buffer B was as follows: 0% at 0min, 10% at 10min, 30% at 135min, 60% at 140min, 100% at 147min, 0% at 150min.

### Untargeted/data-dependent phosphoproteomics mass spectrometry

Data-dependent acquisition (DDA) settings for MS1 scans: m/z range = 350–2000; resolution = 60,000; automatic gain control (AGC) target = 3×10^6^; maximum injection time (maxIT)=50 ms. Highly abundant ions accumulated within 3 seconds cycle time were isolated and fragmented by higher energy collision dissociation (HCD) as follows: resolution= 60,000; AGC target= 1×10^5^; maxIT= 250 ms; isolation width= 0.4 m/z, collision energy (CE)= 33%, dynamic exclusion: if precursor ion is selected 2 times in a 30s window, then exclude for 120s, mass tolerance = 5ppm. IP supernatant was diluted 1:1000 (v:v), loaded on to a pre-column (ID=100µm) packed with 10µm C18 beads, separated using 150 minutes LC gradient, and analyzed on a Q Exactive Plus mass spectrometer in positive polarity mode with MS1 settings: scan range = 350 – 2000m/z, resolution = 70,000, AGC target = 3×10^6^, maxIT=50ms. The top 10 most abundant ions were isolated and fragmented using MS/MS settings: resolution = 35,000, AGC target = 1×10^5^, maxIT=150ms, isolation window = 0.4m/z, NCE=29%, dynamic exclusion=30sec.

### Targeted phosphoproteomics

Precursor ions for the inclusion list were generated by combining targets identified in DDA analysis and enzyme phosphorylation site extracted from PSP for structural annotation. Curated phosphosites were mapped to both human and mouse isoforms of the respective enzyme, and sequences were generated for tryptic peptides of mouse proteome with up to 2 missing cleavages. Inclusion list was generated for each peptide for charge states 2 – 6 with modifications containing TMTpro and methionine oxidation, and allowing for multi-site phosphorylation^143^. Inclusion list was filtered to remove any peptides with length less than 6 or more than 40 amino acids. Additionally, any precursors with charge state [4, 5, 6] and m/z < 400 or charge state [2, 3] and m/z > 1600 were eliminated since they were unlikely to be detected with our LC-MS setup. The inclusion list was then used with the following acquisition settings for an unscheduled targeted analysis: MS1 scan range = 350–1800 m/z; resolution = 120,000; Normalized AGC target = 300%; maxIT=50 ms. The top 30 ions that were in the inclusion list were isolated and fragmented by HCD as follows: mass tolerance = 3ppm, resolution= 120,000; Normalized AGC target = 1000%; maxIT= 247 ms; isolation width= 0.4 m/z, CE= 33%. LC gradient and crude protein analysis were done as stated in the untargeted/data-dependent analysis.

### Phosphoproteomics data analysis

Raw mass spectrometry files were searched in Proteome Discoverer 3.0 powered by Mascot 2.8 using Swissprot for DDA analysis or custom database of tryptic peptides monitored by oru targeted analysis. Searched files were filtered for peptide spectral match (PSM) using the following criteria: Δm/z (ppm) = [-10 – 10] for untargeted and [-5– 5] for targeted; Expectation Score < 0.05; search engine rank = 1; and ions score of >15; ptmRS probability >50%. For each phosphopeptide, TMT intensities of all matching PSMs were summed, and variability across TMT channels was corrected by normalizing to the mean of summed peptides from the crude supernatant. For each phosphopeptide, the summed PSMs were divided by the corresponding value in the bridge sample to normalize across multiple MS runs and generate bridge centered peptide abundance. Following data normalization, we performed hierarchical clustering, principal components analysis (PCA), and sample correlation analysis using Python 3.9 (packages used: Pandas, Numpy, scipy, sklearn, gseapy, matplotlib, and seaborn). Reported multiple hypothesis corrections were done using Benjamini-Hochberg method on p-values obtained from a two-sided Student’s t-test performed on male and female samples separately. All data analysis/visualization was performed using Python 3.9 and GraphPad Prism 10.

### Tissue Metabolite extraction

Polar metabolites were extracted from pulverized mouse liver tissues in 800uL of 60% methanol solution and vortexed for 10min at 4^°^C. Then 500uL of pre-chilled chloroform was added and samples were vortexed for 10min at 4^°^C, followed by centrifugation at 21000xg for 10min at 4^°^C. The top-layer (methanol containing polar metabolites) was transferred to clean 1.7mL tubes, and the interphase (containing protein) was collected in a separate tube for protein quantitation via BCA. Polar metabolites were dried down under a nitrogen drier and stored at -80^°^C before LC-MS analysis. Samples were normalized during resuspension to the total protein content obtained from the BCA of the interphase layer. Briefly, samples were resuspended by vortexing for 10 minutes at 4^°^C in HPLC-grade water containing internal standards: U-^13^C labeled yeast metabolite extract at 1:20 (v:v) and U-^13^C, ^15^N labeled amino acids at 500nM final concentration.

### Polar LC-MS Metabolomics

For each sample 2uL of resuspended polar metabolites were injected on to SeQuant ZIC-pHILIC 5 μm 150 × 2.1 mm analytical column in line with a SeQuant ZIC-pHILIC 2.1 × 20 mm guard column. Analytes were separated using a Vanquish Neo UPLC system coupled to a Q Exactive Orbitrap mass spectrometer. Chromatography was performed at 0.150mL/min flow rate with linear gradient of 80 – 20% of buffer B (100% Acetonitrile) for 20min, followed by 20 – 80% buffer B at 20.5min and held at 80% buffer B for the last 8min, totaling 28min in a 20mM ammonium carbonate in 0.1% ammonium hydroxide buffer A. Data was acquired on a Q Exactive Orbitrap mass spectrometer in full-scan polarity-switching mode with MS1 settings: range = 70 – 1000m/z, resolution = 70,000, AGC = 1 x 10^6^, and maximum IT = 20ms. To maximize detection of nucleotides, an additional scan was performed in negative mode with MS1 settings: range = 200 – 700m/z, resolution = 70,000, AGC = 1 x 10^6^, and maximum IT = 80ms. Relative quantitation of metabolites was performed with TraceFinder 5.2 a 4 mmu mass tolerance and referencing retention times from both in-house library of small molecule standards as well as in-run U-^13^C labeled yeast metabolites and U-^13^C, ^15^N labeled amino acids. Mean centered raw peak areas were used to perform hierarchical clustering, PCA, and spearman correlation analysis in Python 3.9 using packages: pandas, numpy, scikit-learn, scipy, matplotlib, and seaborn.

### Plasmids and cloning reagents

SuperPiggyBac Transposase purchased from SystemBio was linearized with SmaI and HindIII, then cloned into pUC19 vector linearized using SmaI and HindIII. HA-KRAB-dCas9 and PB_rtTA_BsmBi vectors were purchased from Addgene^85^, and the Neomycin selection marker in PB_rtTA_Bsmbi was switched out for Blasticidin. The single guide RNA (sgRNA) sequences were generated using GPP web portal tool from the Broad Institute^144–146^ with Human GRCh38 for SpyoCas9 (NGG), where the top 4 sequences were used to generate plasmids in PB_rtTA_BsmBi vector. Briefly, gene specific sequences were individually ligated into PB_rtTA_BsmBi vector linearized using BsmBI-v2, generating 4 sgRNA plasmids. Plasmids for wildtype enzyme open reading frames (ORFs) were purchased from Addgene or DNASU plasmid repository: pDONOR221_GSTP1, pDonor223_IDH1, pDONOR221_UMPS. To generate phosphomimic or phosphodeficient mutants, site-directed mutagenesis of wildtype ORFs was done using Q5 site-directed mutagenesis kit with primers generated using NEBaseChanger. All ORFs were gateway cloned into pCW57.1 designation vector using LR Clonase II. For *in vitro* assays using recombinant GSTP1, ORFs were cloned into pHAGE-CMV-FLAG^147^ using LR Clonase II. All oligonucleotide sequences used for cloning are provided in Table S15.

### Cell Culture

A549 and HEK293T cell lines was obtained from the American Type Culture Collection (ATCC) and used within 10 passages after receipt from ATCC to ensure their identities. Cells were cultured at 37^°^C in a humidified incubator with 5% CO2. Cells were passaged with 0.05% Trypsin/0.53mM EDTA in Sodium Bicarbonate, and maintained in Dulbecco’s Modified Eagle Medium (DMEM) supplemented with 10% tetracycline-negative fetal bovine serum (FBS).

### Generation of CRISPRi Cell lines

Parental A549 cells were plated 24 hours before transfection. The following day cells were transiently co-transfected with SuperPiggyBac Transposase, HA-KRAB-dCas9 and pool of 4 sgRNA vectors at a ratio of 2:1:1 (weight in µg) of DNA using 20µg of PEI-max transfection reagent for 36 hours. Polyclonal lines were generated via selection with 5µg/mL Blasticidin and 300µg/mL of Hygromycin B for 2 passages, then maintained in 0.5µg/mL Blasticidin and 50µg/mL Hygromycin B.

To express inducible rescue or control ORFs in pCW57.1 backbone, lentivirus was generated using PsPax2 and PMD2G packaging vectors in HEK293Ts. Briefly, HEK293T cells were plated in 10cm plates for 24 hours, then transfected with the respective ORF vectors, PsPax2, and PMD2G at a ratio of 10:10:5 µg of DNA using PEI-max. After 16 hours of transfection media was replaced to DMEM with 20% FBS, and lentivirus was collected 48 hours after media change. A549 cell lines stably expressing sgRNAs and inducible HA-KRAB-dCas9 were infected with the respective inducible rescue construct or control mCherry lentivirus supplemented with Polybrene. After 36 hours of infection, media was changed to 5µg/mL Puromycin selection for 2 passages, and then maintained in 0.5µg/mL Puromycin, 0.5µg/mL Blasticidin, and 50µg/mL Hygromycin B.

### Western blot

A549 CRISPRi-rescue cells were plated at 5000 cells/mL in 6-well plates, and 24 hours later treated with 500ng/mL of doxycycline (Dox) replenished daily for 96 hours total. All samples were lysed in RIPA (10% glycerol, 50mM Tris-HCl, 100mM NaCl, 2mM EDTA, 0.1% SDS, 1% Nonidet P-40, 0.2% Sodium Deoxycholate) supplemented with Halt™ Protease and Phosphatase Inhibitor Cocktail and Benzonase. Lysis was done on ice for 30 minutes and lysates were centrifuged for 15 minutes at 21,000xg at 4^°^C. Protein amount was normalized via BCA, and samples were denatured in NuPAGE LDS buffer with 1mM DTT. Antibodies used for western blots are as follows: GSTP1, IDH1, UMPS, Vinculin, HA, PCNA.

### Generation of recombinant GSTP1 & *in vitro* phosphorylation

HEK293T cells were plated in 10cm plates for 24 hours, then transiently transfected with 25µg of the respective Flag-GSTP1 variant or Flag-mCherry control for 24 hours. Cells were lysed in IP buffer (0.3% NP-40, 50mM Tris-HCl, 150mM NaCl) and sonicated at 10% amplitude for 3 – 5 seconds at 4^°^C, then lysates were cleared via centrifugation at 21,000xg for 15 minutes at 4^°^C. Then samples were divided into two sets per variant to generate either control or hyperphosphorylated lysates using recombinant SRC. Briefly, 0.5µg of recombinant SRC kinase was mixed with 10x kinase buffer (200mM HEPES pH 7.5, 100mM DTT, 1mM EGTA, 1mM ATP, 100mM MgCl2, 100mM MnCl2) and added at a 1:10 (v/v) ratio to the +SRC lysate, and for control reactions kinase buffer without SRC was used. Lysates were incubated for 30min at room temperature, then immediately placed on ice for immunoprecipitation (IP) using magnetic FlagM2 beads. Flag IP was done overnight at 4^°^C, after which beads were washed 3x with wash buffer (50mM Tris HCl, 150mM NaCl, pH 7.4), then 1x with molecular grade water. Proteins were then eluted twice in 0.1M Glycine HCl at pH 3.5 and neutralized by adding 1:10 (v:v) of neutralization buffer (1.5M NaCl, 0.5M Tris HCl, pH 7.4). Protein concentration was determined using BCA assay, and expression was confirmed by running 1µg of protein denatured in Laemmli-SDS reducing buffer containing 2-mercaptoethanol at 95^°^C for 5 minutes, then separated via gel electrophoresis and visualized with SimplyBlue Safestain.

### GSTP1 activity assay

GSTP1 enzyme assays were done using commercial GST assay kit and per manufactures protocol for both cell based and in vitro assays. Briefly, cell-based assays were done by plating A549 CRISPRi-rescue cells at 5000 cells/mL in 6-well plates, then 24 hours later 500ng/mL Dox was added daily for 96 hours total. Cells were then washed with PBS and lysed in sample buffer supplied with the GST kit. For assays using recombinant Flag-GSTP1 variants, 1µg of protein was diluted in sample buffer. Assays were run in 384-well plate with a final reaction volume of 20µL per well (10µL sample, 2µL glutathione, 8µL assay buffer), and each biological replicate was assayed in technical triplets. Statistical analysis was performed on the means of technical replicates across 4 biological replicates using Two-way ANOVA with multiple hypothesis correction (Dunnett) in GraphPad Prism10 where statistical significance was assigned for p< 0.05.

### U-^13^C5-Glutamine tracing

A549 CRISPRi-rescue cells plated as stated above, and after 96 hours of dox induction, cells were washed and given media with 2mM of unlabeled glutamine or U-^13^C5-Glutamine. Tracing was done for 1 & 2 hours for IDH1 or 12 & 24 hours for UMPS. Afterwards, cells were washed with blood bank saline, and metabolites were extracted using 80% methanol in 20% HPLC grade water supplemented with 8µg/mL Norvaline as an internal standard. Samples were vortexed for 10 minutes at 4^°^C then centrifuged at 21,000xg for 10 minutes at 4^°^C, where the supernatant was collected and dried under nitrogen gas.

### GC-MS metabolomics analysis of U-^13^C5-Glutamine tracing

Dried polar metabolites were derivatized by incubating in 16µL Methoxamine (MOX) Reagent at 37^°^C for 1 hour, followed by incubation in 20µL N-tert-Butyldimethylsilyl-N-methyltrifluoroacetamide with 1% tert-Butyldimethylchlorosilane (TBDMS) at 60^°^C for 1 hour. Derivatized samples were then quickly centrifuged, and analyzed using DB-35MS column (30 m x 0.25 mm i.d. x 0.25 μm) in an Agilent 7890 gas chromatograph (GC) coupled to an Agilent 5975C mass spectrometer (MS). 1µL of sample was introduced in split mode 1:1 with a helium carrier gas at a constant flow rate of 1.2mL/min. Post-injection, the GC oven was held at 100^°^C for 1 minute, then gradually raised to 105^°^C at a rate of 2.5^°^C/min and held at 105^°^C for 2 minute. The temperature was then increased to 250^°^C at 3.5^°^C/min and finally ramped at 320^°^C at a rate of 20^°^C/min. The MS system was operated with electron impact ionization of 70eV, where the source and quadrupole were kept at 230^°^C and 150^°^C, respectively. Raw GC-MS files were converted to mZML using MS Convert v.3.0^148^, and peak areas were obtained using El-Mavin v0.11.0^149^ followed by natural isotope abundance correction using IsoCorrectorGui ^150,151^. All ion counts were normalized to the internal standard, norvaline, and protein quantification of samples.

### LC-MS metabolomics analysis of U-^13^C5-Glutamine tracing

Dried polar metabolites were resuspended in HPLC grade water, and 5µL of resuspended polar metabolites were injected on to SeQuant ZIC-pHILIC 5 μm 150 × 2.1 mm analytical column. Analytes were separated using an Ultimate 3000 UPLC system coupled to a Q Exactive Orbitrap mass spectrometer. Chromatography was performed at 0.150mL/min flow rate with linear gradient of 80 – 20% of buffer B (100% Acetonitrile) for 20min, followed by 20 – 80% buffer B at 20.5min and held at 80% buffer B for the last 8min, totaling 28min in a 20mM ammonium carbonate with 0.1% ammonium hydroxide buffer A. Mass spectrometry data acquisition was done in full-scan polarity-switching mode with MS1 settings: range = 70 – 1050m/z, resolution = 70,000, AGC = 1 x 10^6^, and maxIT = 80ms. Additional scans were done to detect nucleotides in negative mode with MS1 settings: range = 200 – 700m/z, resolution = 70,000, AGC = 1 x 10^6^, and maximum IT = 80ms. Relative quantitation of metabolites was performed with TraceFinder 5.2 with 5 ppm mass tolerance and referencing retention times from in-house library of chemical standards. Natural isotope abundance correction was done using IsoCorrectorGui ^150,151^. All ion counts were normalized to the internal standard, norvaline, and protein quantification of samples.

### Partial least squares regression (PLSR) analysis

Bridge centered phosphoproteomics and mean centered metabolomics datasets were used as input for PLSR analysis. PLSR model was run for each metabolite, where pY dataset was used as the X-matrix, and a metabolite as the Y-matrix. In order to capture the variance within both omics datasets, we performed feature selection to filter for the top 50% of pY site with the highest variance. For the first cohort of in vivo studies using HFD and NCD samples, PLSR was performed using k-fold cross-validation with n_splits = 7, such that the sample set was divided into 7 groups. Then the PLSR model was trained via leave-one-out cross-validation, where 6 groups were used for training to predict metabolite-pY correlation for the remaining group. For BHA studies, only leave-one-out cross-validation was used due to smaller samples size.

Prediction scores (Q^2^) cutoff of 0.4 or more was used to determine valid models without overfitting, and a variable of importance in projection (VIP) score of 1 or above was used to define pY sites that predict the metabolite with a valid model prediction score. Analysis was performed using Python 3.9 with the following packages: numpy, pandas, scipy, and sklearn. See data and code availability section to access results and script.

### Data and code availability

Code to evaluate quality of PDB structures and annotate phosphosites is available in the following Github repository: github.com/phosphoTig/metaPhosphosites.

All code used to process and visualize searched phosphoproteomics and metabolomics analysis is available in the Github repository: github.com/phosphoTig/omicsAnalysis.

The mass spectrometry proteomics raw files and search results have been deposited to the ProteomeXchange Consortium via the PRIDE^152,153^ partner repository with the dataset identifier PXD054497 and 10.6019/PXD054497.

Polar metabolomics data for this study is available at the NIH Common Fund’s National Metabolomics Data Repository (NMDR) website, the Metabolomics Workbench^154^, where it has been assigned Study ID ST003375. The data can be accessed directly via its Project DOI: https://dx.doi.org/10.21228/M82V51.

## Acknowledgements

We would like to thank Dr. Marino Convertino for invaluable scientific discussion of computational structural analysis and life guidance. We would also like to thank members of the White lab and the Vander Heiden lab for scientific and technical exchange of ideas. We acknowledge the MIT Undergraduate Research Opportunities Program (UROP) for supporting SC, AXL, RM, and FNV. This work was supported in part by the Koch Institute Support (core) Grant 5P30-CA014051 from the National Cancer Institute. We thank the Koch Institute’s Robert A. Swanson (1969) Biotechnology Center for technical/resource support; specifically Flow cytometry, High Throughput Sciences, and Bioinformatics cores. Public access to metabolomics data via Metabolomics Workbench is supported by NIH grant U2C-DK119886 and OT2-OD030544 grants.

## Author Contributions

This work was designed by TYT and FMW. Experiments were carried out by TYT, AL, NJK, SET, CTF, PV, YC, and JCD. Computational analyses were performed by TYT, SC, RM, and FV. Manuscript was written by TYT and FMW. Manuscript was reviewed and edited by TYT, FMW, CTF, SET, JCD, ALH, AT, MGVH, JBS, and RJD.

| Reagent or Resource | Source | Identifier/part number |
| --- | --- | --- |
| **Mouse Work** |  |  |
| C57BL6/J | The Jackson Laboratories | 000664 |
| High fat diet (HFD) | Bio-Serv | S3282 |
| Normal chow diet | Bio-Serv | F4031 |
| HFD + 1.5% 2(3)-tert-butyl-4 hydroxyanisole (BHA) | Bio-Serv | S7958 |
| **Phosphoproteomics** |  |  |
| 1260-Infinity LC Systems | Agilent |  |
| Orbitrap Exploris™ 480 Mass Spectrometer | Thermo Fisher Scientific |  |
| Orbitrap Q-Exactive Plus | Thermo Fisher Scientific |  |
| Syringe Pump | Harvard Apparatus |  |
| Tissue Homogenizer | VWR |  |
| Speed Vac | Thermo Fisher Scientific |  |
| Infinite 200 Pro Microplate reader | Tecan |  |
| 3µm C18, ODS-AQ, 12nm | YMC | AQ12S03 |
| 10µm C18, ODS-AQ, 12nm | YMC | AA12S11 |
| 200µm ID fused silica capillary | Molex, LLC-Polymicro Technologies | TSP200350 |
| 100µm ID fused silica capillary | Molex, LLC-Polymicro Technologies | TSP100375 |
| 50µm ID fused silica capillary | Molex, LLC-Polymicro Technologies | TSP050375 |
| Urea | Millipore Sigma | U51288 |
| Pierce™ BCA Assay Kit | Thermo Fisher Scientific | 23225 |
| DTT | Millipore Sigma | D9779 |
| Iodoacetamide | Millipore Sigma | I1149 |
| Ammonium Acetate | Millipore Sigma | A7330 |
| Sequencing grade Trypsin | Promega | V511 |
| Glacial Acetic Acid | Thomas Scientific | 216175 |
| SepPak Light C-18 cartridges | Waters Corporation | WAT023501 |
| HEPES | Corning | 25-060-CI |
| TMT 16plex Reagent | Thermo Fisher Scientific | A44520 |
| High-Select™ Fe-NTA Phosphopeptide Enrichment Kit | Thermo Fisher Scientific | A32992 |
| 50% Hydroxylamine | Thermo Fisher Scientific | 90115 |
| PT66 | Millipore Sigma | P3300 |
| Super 4G10 anti-pY antibody | Produced in house<sup>141,142</sup> |  |
| Protein G Plus-Agarose Suspension | Millipore Sigma | IP04 |
| Acetonitrile | Millipore Sigma | 34998 |
| Acetic Acid (LC-MS) | Millipore Sigma | 338826 |
| Trifluoroacetic Acid | Millipore Sigma | 299537 |
| **Metabolomics** |  |  |
| Nitrogen Drier | VWR |  |
| Vanquish Neo UPLC System | Thermo Fisher Scientific |  |
| Ultimate 3000 UPLC System | Thermo Fisher Scientific |  |
| Agilent 7890 Gas Chromatograph | Agilent |  |
| Agilent 5975C Mass Spectrometer | Agilent |  |
| U-<sup>13</sup>C yeast metabolite extract | Cambridge Isotope Laboratories | ISO1 |
| U-<sup>13</sup>C, <sup>15</sup>N amino acid mix (17 AA) | Cambridge Isotope Laboratories | MSK-A2-1.2 |
| Blood bank saline | Azer Scientific | ES12444G |
| Norvaline | Millipore Sigma | N7627 |
| SeQuant ZIC-pHILIC 5 µm 150 × 2.1 mm analytical column | Millipore Sigma | 1504600001 |
| SeQuant ZIC-pHILIC 20 × 2.1 mm guard column | Millipore Sigma | 150438001 |
| U-<sup>13</sup>C<sub>5</sub>-Glutamine | Cambridge Isotope Laboratories | CLM-1822-H |
| Methoxamine (MOX) | Thermo Fisher Scientific | TS45950 |
| Tert-Butyldimethylsilyl-N-methyltrifluoroacetamide with 1% tert-Butyldimethylchlorosilane (TBDMS) | Millipore Sigma | 375934 |
| DB-35MS column (30m x 0.25 mm i.d. x 0.25µm) | Agilent J&W Scientific | 122-3832UI |
| **Plasmids and cloning** |  |  |
| SuperPiggyBac Transposase | SystemBio | PB210PA-1 |
| SmaI | New England Biolabs | R0141 |
| HindIII | New England Biolabs | R0104 |
| BsmBI-v2 | New England Biolabs | R0739 |
| pUC19 backbone | Thermo Fisher Scientific | SD0061 |
| PsPax2 | Addgene | 12259 |
| PMD2G | Addgene | 12260 |
| pCW57.1 | Addgene | 41393 |
| HA-KRAB-dCas9 | Addgene | 126030 |
| PB_rtTA_BsmBI | Addgene | 126028 |
| IDH1 (pDONR223) | Addgene | 82133 |
| GSTP1 (pDONR221) | DNASU | HsCD0004479 |
| UMPS (pDONR221) | DNASU | HsCD00296310 |
| Q5 site-directed mutagenesis kit | New England Biolabs | E0552S |
| LR clonase II | Thermo Fisher Scientific | 11791-020 |
| pHAGE-CMV-FLAG | Tamir et al. 2020, JCS<sup>147</sup> |  |
| mCherry (pDONR223) | Tamir et al. 2020, JCS<sup>147</sup> |  |
| **Cell culture** |  |  |
| Dulbecco's Modified Eagle Medium (DMEM) | Corning | 10-013-CV |
| DMEM without glutamine, sodium pyruvate, and phenol red | Gibco | 31053-028 |
| Sodium pyruvate | Corning | 25-000-CI |
| tetracycline-negative fetal bovine serum (FBS) | GeminiBio | 100-800-500 |
| Dialyzed FBS | Gibco | 26400-044 |
| A549 | ATCC | CCL-185 |
| HEK293T | ATCC | CCL-3216 |
| 0.05% Trypsin/0.53mM EDTA in Sodium Bicarbonate | Corning | 25-052-CI |
| Doxycycline | Millipore Sigma | D9891 |
| L-Glutamine | Corning | 25-005-CI |
| Blasticidin | Gibco | A1113903 |
| Hygromycin B | Thermo Fisher Scientific | 10687010 |
| Puromycin | Thermo Fisher Scientific | A1113803 |
| Polybrene | Millipore Sigma | TR-1003-G |
| PEI-Max | Polysciences Inc | 24765-1 |
| Opti-MEM | Thermo Fisher Scientific | 31985062 |
| **Protein gels & Western blots** |  |  |
| Halt™ Protease and Phosphatase Inhibitor Cocktail | Thermo Fisher Scientific | 78445 |
| Benzonase | Millipore Sigma | E1014 |
| NuPAGE LDS buffer | Thermo Fisher Scientific | NP0007 |
| Laemmli-SDS reducing buffer | Boston BioProducts | BP-111R |
| Precision Plus Protein Dual Color Standards | Bio-rad | 1610374 |
| 12+2 well BioRad CriterionX Bis-Tris Gel (4-12%) | Bio-rad | 3450123 |
| 26 well BioRad CriterionX Bis-Tris Gel (4-12%) | Bio-rad | 3450125 |
| Nitrocellulose Membrane | Bio-rad | 1620112 |
| SuperSignal™ West Pico PLUS Chemiluminescent Substrate | Thermo Fisher Scientific | 34577 |
| SimplyBlue Safestain | Thermo Fisher Scientific | LC6060 |
| **Antibodies** |  |  |
| GSTP1 | Cell Signaling Technology | 3369 |
| IDH1 | Cell Signaling Technology | 8137 |
| HA | Cell Signaling Technology | 2367 |
| PCNA | Cell Signaling Technology | 2586 |
| UMPS | Bethyl/Fortis Life Sciences | A304-258A-M |
| Vinculin | Millipore Sigma | V4889 |
| Goat anti-Mouse secondary | Jackson ImmunoResearch Laboratories | 115-035-0174 |
| Mouse anti-Rabbit secondary | Jackson ImmunoResearch Laboratories | 211-032-171 |
| **In Vitro GSTP1 phosphorylation** |  |  |
| ProQinase™ SRC (HIS-tag) | Reaction Biology | 0260-0000-1 |
| FlagM2 magnetic beads | Millipore Sigma | M8823 |
| Molecular Grade Water | Corning | 46-000-CI |
| EGTA (0.5M) | Boston BioProducts | BM-721 |
| ATP (100mM) | VWR | 10191-288 |
| MgCl<sub>2</sub> (1M) | Thermo Fisher Scientific | AM9530G |
| MnCl<sub>2</sub> (1M) | Millipore Sigma | M1787 |
| Glycine (0.2M, pH=3.0) | Boston BioProducts | BB-95-LT |
| Tris-HCl (1M, pH=7.4) | Boston BioProducts | BBT-74 |
| GST assay kit | Cayman | 703302 |
| 384 well plates | Millipore Sigma | CLS3702-100EA |
| **Oligonucleotides** |  |  |
| See Table S15 | This paper |  |
| **Databases** |  |  |
| PhosphoSitePlus (PSP) | Cell Signaling Technology | Version: December 2023 |
| Kyoto Encyclopedia of Genes and Genomes (KEGG) Pathway |  |  |
| UniprotKB |  | Version: 2023 |
| RCSB PDB |  |  |
| PDBePISA |  |  |
| Swissprot |  |  |
| Mouse phosphopeptide database | This paper | PRIDE: PXD054497 |
| **Deposited Data** |  |  |
| Phosphoproteomics data | This paper | PRIDE: PXD054497 |
| Steady state metabolomics data | This paper | Metabolomics Workbench: ST003375 |
| **Software and Algorithm** |  |  |
| GraphPad Prism 10.3 |  |  |
| Pymol Version 3.0 | Schrödinger, LLC |  |
| Python 3.9 |  |  |
| TraceFinder 5.2 | Thermo Fisher Scientific |  |
| ProteomeDiscoverer 3.0 | Thermo Fisher Scientific |  |
| Mascot 2.8 | Matrix Science |  |
| Structural annotation | This paper | Git-hub:<br>phosphoTig/metaPhosphosites |
| Omics data analysis | This paper | Git-hub:<br>phosphoTig/omicsAnalysis |

## Supplemental information index

**Document S1:** Figures S1 – S7 and Tables S4, S11 – 13, S15

Table S1: Excel file containing enrichment analysis related to Figures 1C, 1D

Table S2: PDB quality assessment related to Figure 2A (available upon request)

Table S3: Phosphosite-domain distances related to Figures 2B - 2C (available upon request)

Table S4: Hypergeometric distribution test results related to Figure 2D

Table S5: PISA results of interface phosphosites related to Figures 2E, 2F

Table S6: Steady state polar metabolomics dataset related to Figures 3B – 3I and 5B – 5J

Table S7: pY phosphoproteomics dataset related to Figures 3H-3M

Table S8: PLSR analysis results related to Figure 4B

Table S9: pY- Domain proximity of predictive sites related to Figure 4C (available upon request)

Table S10: PLSR analysis results on BHA treated samples related to Figure 5K **Table S11:** GSTP1 activity assay on CRISPRi-rescue A549 Cells related to Figure 6B **Table S12:** GSTP1 activity assay on recombinant GSTP1 protein related to Figure 6D **Table S13:** Western blot quantitation of GSTP1 knockdown related to Figure 6B

Table S14: Summary of U-^13^C5Glutamine tracing for IDH1 and UMPS related to Figure 7

Table S15: Oligonucleotide sequences for CRISPRi and site-directed mutagenesis

---

**Figure S1.**
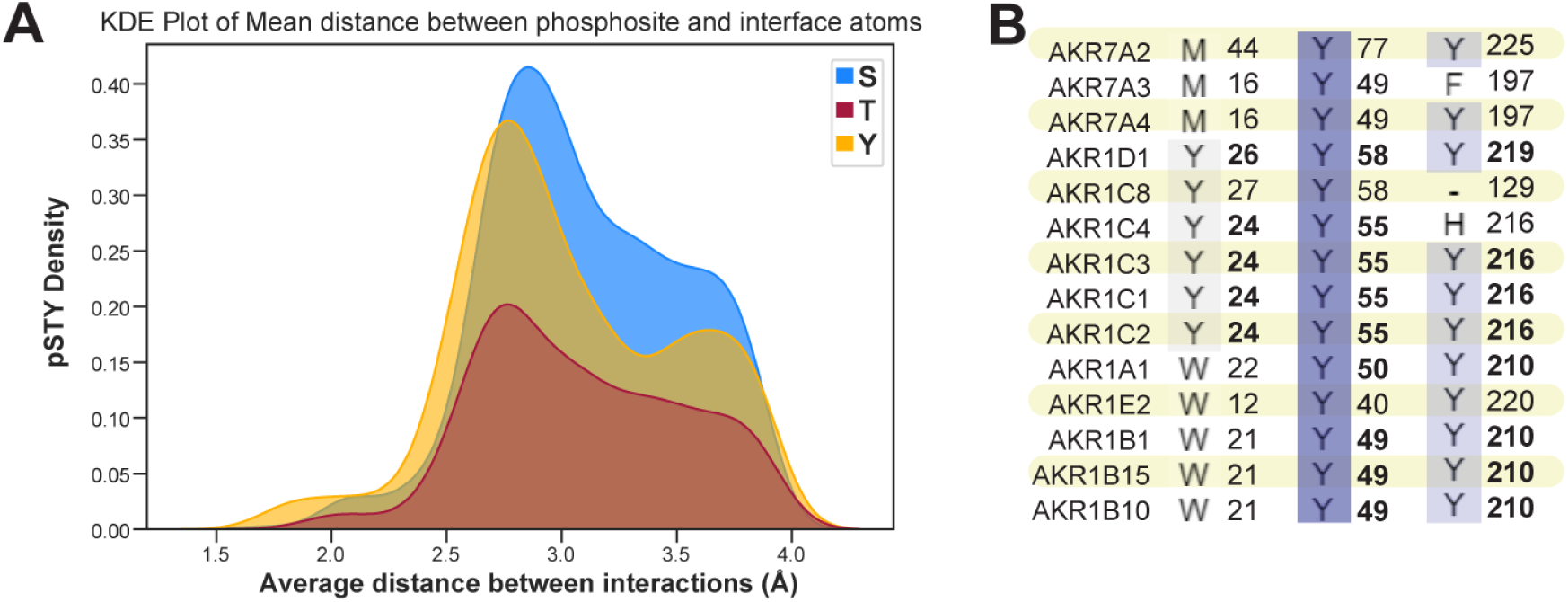
Phosphosites on metabolic enzymes are highly conserved, and the hydroxyl groups interface phosphosites form contacts with residues and ligands at or below a distance of 4Å. (A) Kernel density estimate (KED) plot showing the distribution of distances between phosphosites and interface residues or ligands. **(B)** UniprotKB sequence alignment to illustrate conservation of pY sites within the active site and cofactor binding domain of AKR family of enzymes.

**Figure S2.**
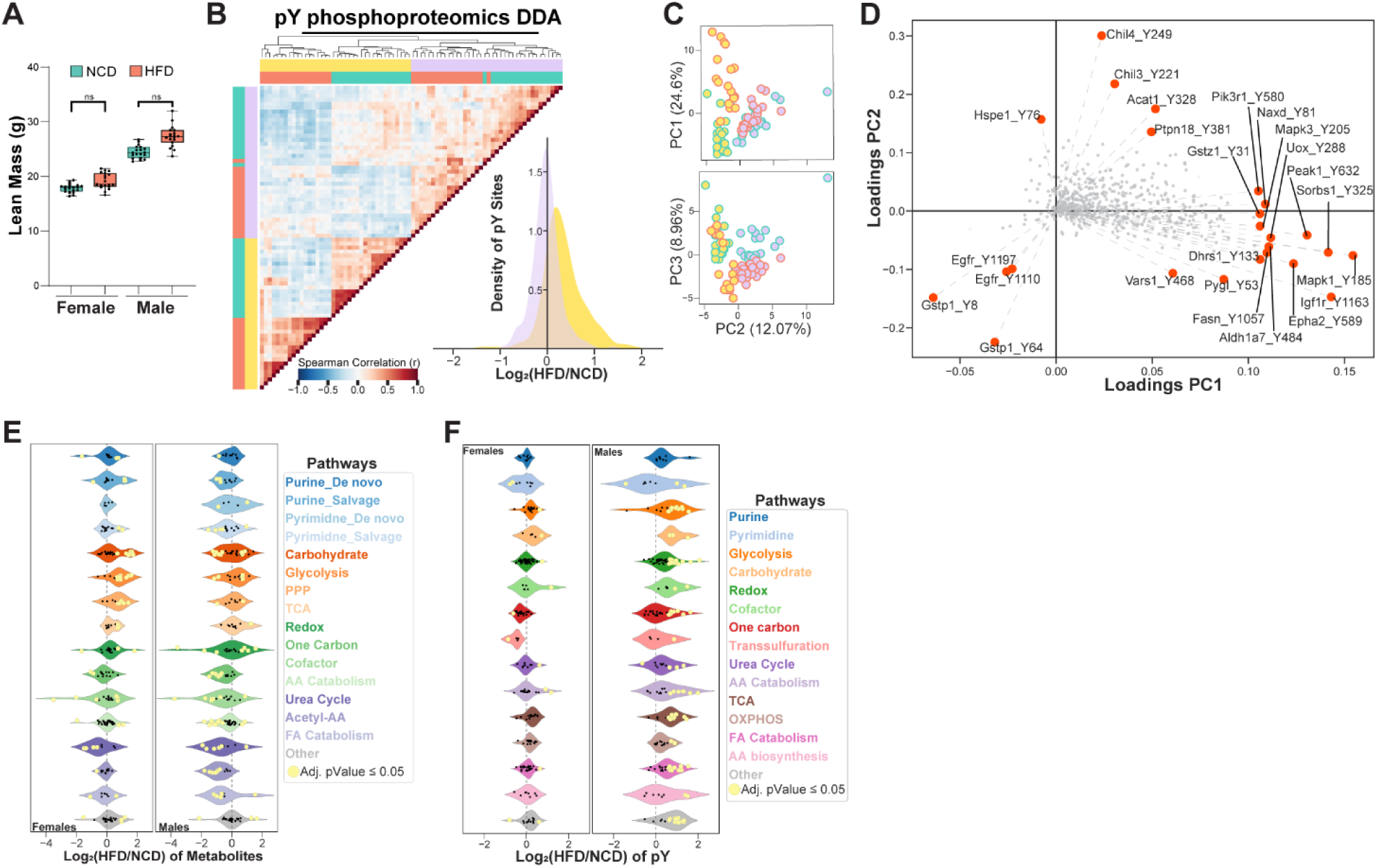
HFD does not significantly alter lean mass; and while untargeted pY phosphoproteomics captures similar trends, HFD-induced changes in metabolic enzyme pY levels were masked by more abundant signaling proteins. (A) Box and whisker plot of lean mass (g), each point represents individual animal. **(B-D)** heatmap of spearman correlation, KDE plot of fold changes, and scatter plots of PCA loadings for untargeted phosphoproteomics analysis of pY sites. **(E-F)** Pathway specific violin plots of metabolites and pY sites. Metabolites or pY sites that were significantly altered are indicated in yellow (adjusted p-value ≤ 0.05). Two-tailed Student’s t-test was performed and multiple hypothesis correction was done using Benjamini- Hochberg method.

**Figure S3.**
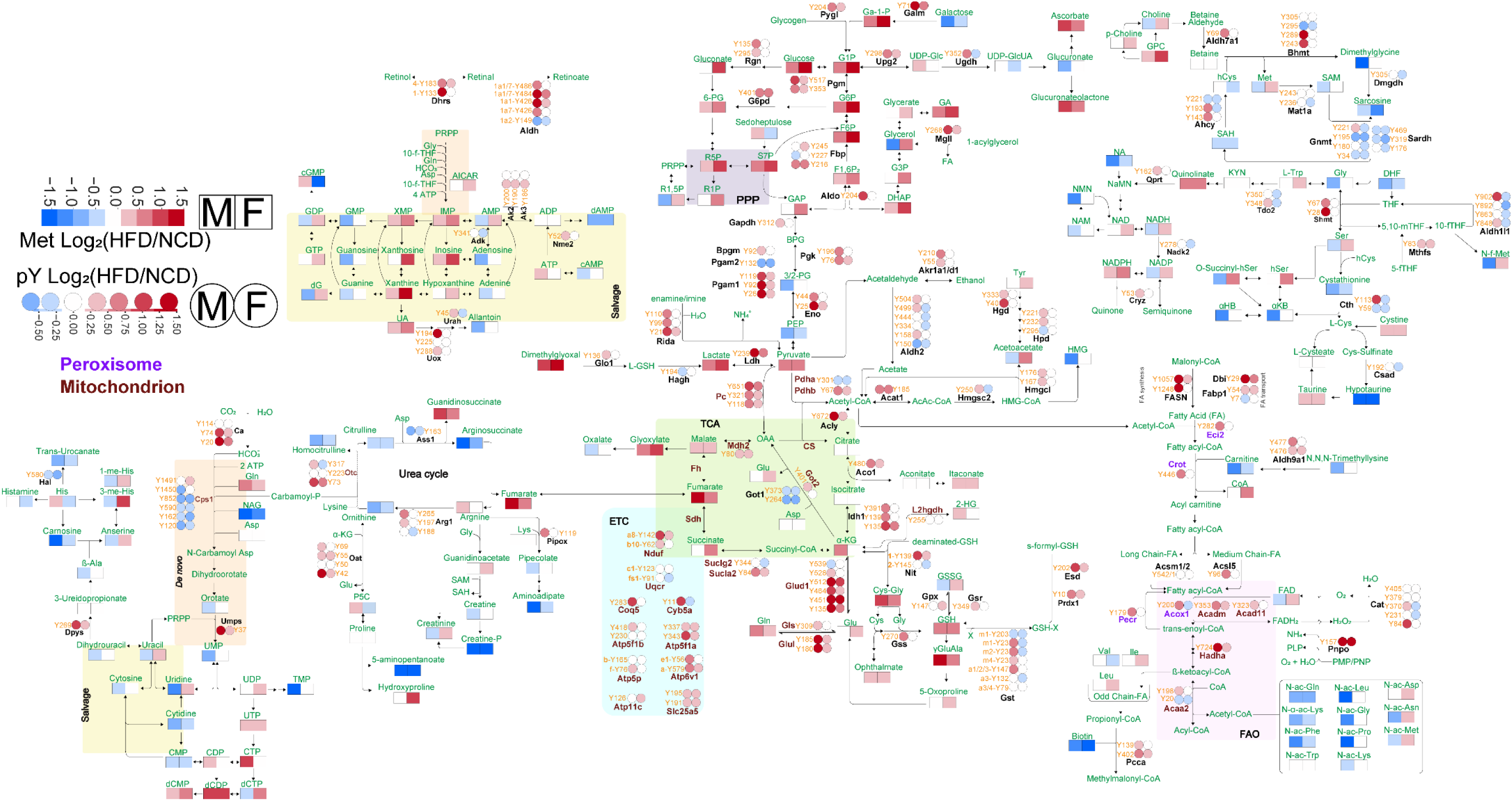
Pathway map of HFD-altered metabolite and pY sites illustrates metabolic reprogramming and signaling input on metabolic enzymes.

**Figure S4.**
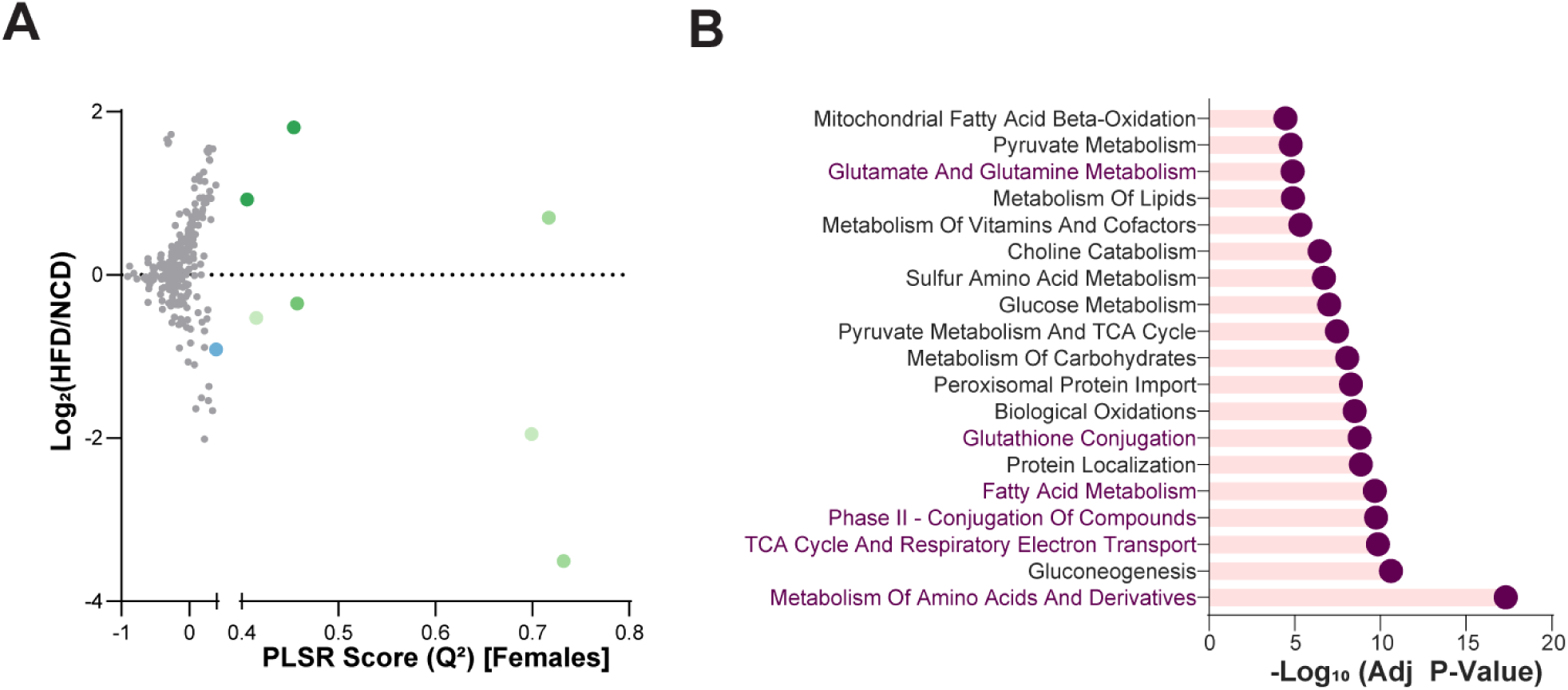
Predictive pY sites from PLSR model are associated with catabolic and oxidative metabolism. (A) Scatter plot representation of PLSR model score (Q^2^) vs Log2(HFD/NCD) for females. Each datapoint represents a metabolite. **(B)** Bar plot of gene level pathway enrichment analysis of enzymes with predictive pY sites, pathways relevant for redox homeostasis and HFD driven phenotype are highlighted in maroon.

**Figure S5.**
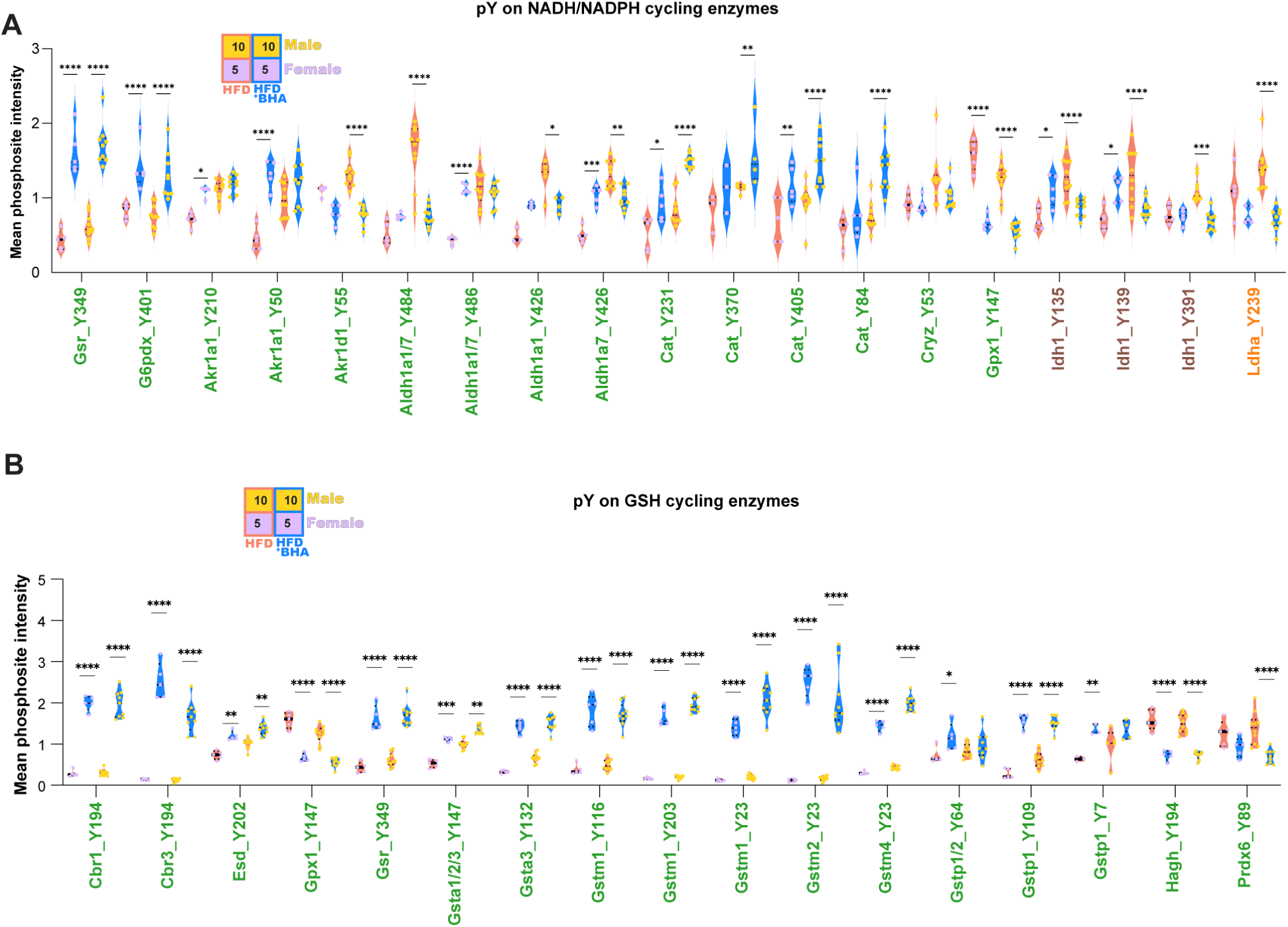
BHA induces differential regulation of pY sites on NADH/NADPH and GSH cycling enzymes. Violin plots for pY sites on enzymes that cycle **(A)** NADH/NADPH and **(B)** GSH, each datapoint represents individual animal. Statistical analysis was done via two-way ANOVA with multiple comparison performed between HFD + BHA and HFD only groups where significance is assigned as follows: * p < 0.05, ** p < 0.01, **** p < 0.001, **** p < 0.0001.

**Figure S6.**
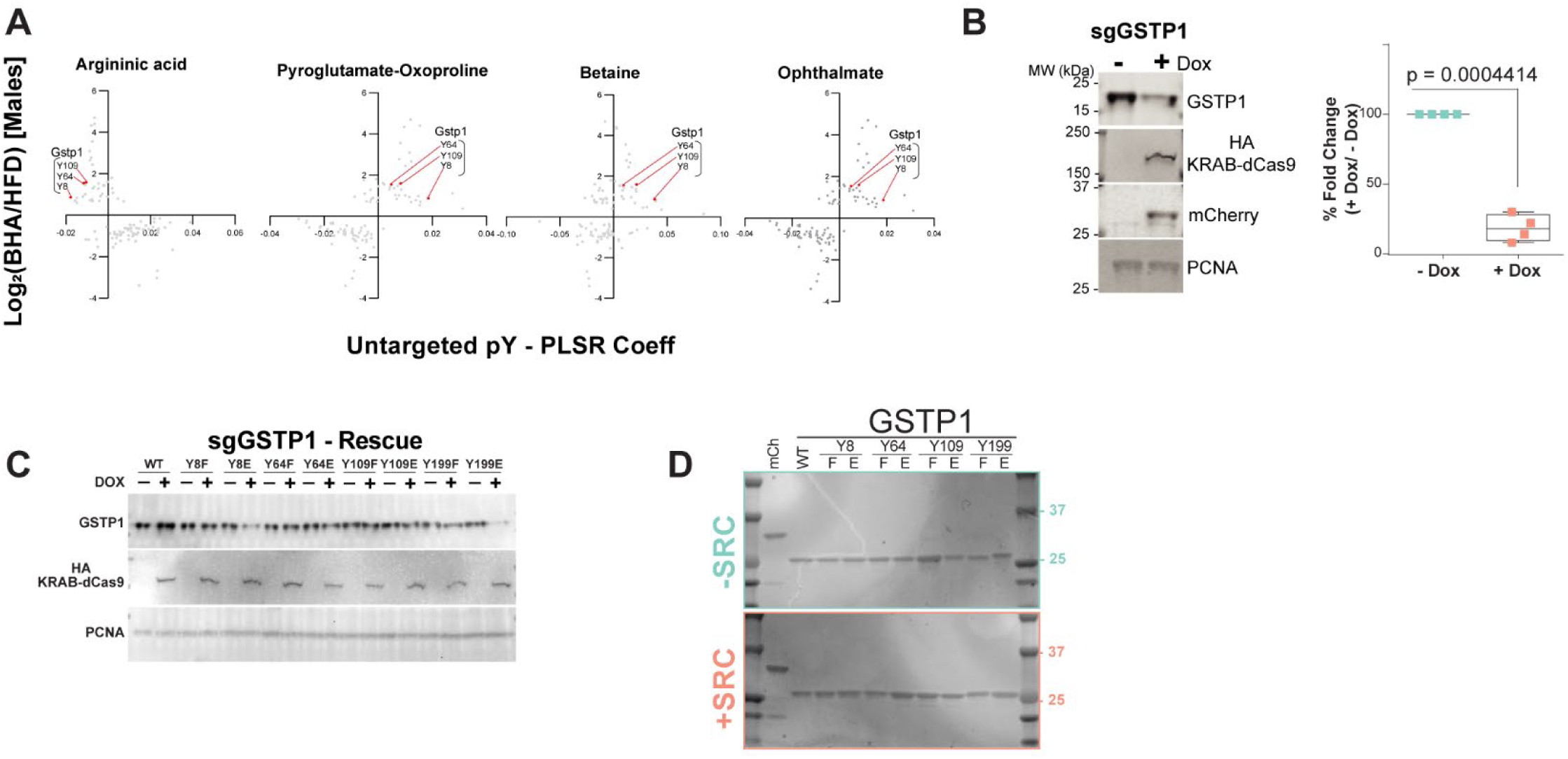
Conserved GSTP1 pY sites predict metabolite alteration in response to BHA. (A) Scatterplots of PLSR analysis for metabolites indicating predictive GSTP1 pY sites (Y8, Y64, Y109) of differential response to BHA for argininic acid, pyroglutamate/oxoproline, betaine, and ophthalmate. Each datapoint represents pY site. **(B)** Sequence similarity matrix heatmap of GST isoforms, and conservation of residues Y8, Y64, and Y109. Analysis generated via UniprotKB. **(C, D)** western blot and quantitation of CRISPRi-mediated knockdown and rescue of human GSTP1 with a pool of 4 single guide RNAs (sgGSTP1) for 96 hours with 500ng/mL of Dox in A549 cells (representative of N=4). Statistical analysis was done using two-tailed, paired t- test. **(E)** Protein gel confirming expression of recombinant GSTP1 phospho-variants used for in vitro assays (representative of N=4).

**Figure S7.**
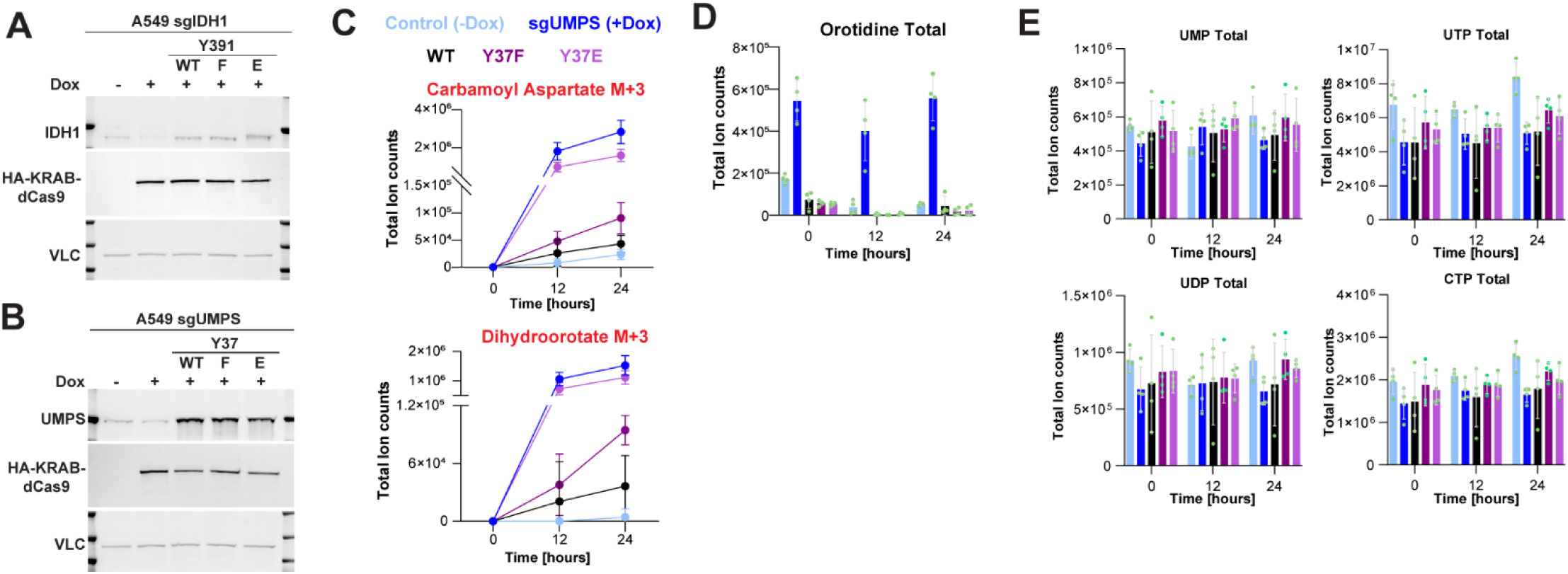
Confirmation of CRISPRi-rescue expression and metabolite changes. (A, B) Western blot validation of CRISPRi-rescue protein expression for IDH1 and UMPS expression. **(C)** Total ion count measurement of M+3, non-dominant species, of labeled carbamoyl aspartate and dihydroorotate (N=4), normalized to Norvaline internal standard and protein content. **(D)** Total ion count of orotidine, first reaction mediated by UMPS, over 0, 12, and 24 hours. **(E)** Total UMP, UDP, UTP, and CTP levels measured from the sum of total ion counts of all observed species.

**Table S4:** Hypergeometric distribution test of phosphosites within 23A of functional domains, associated with. **Figure 2D**.

| DomainType | N (total AA count) | M (total phospho count) | s ( total count w/in 23A) | k (phospho count w/in 23A) | p_value | FDR | reject H0 |
| --- | --- | --- | --- | --- | --- | --- | --- |
| STY_Active Site | 18231 | 2913 | 7977 | 1375 | 2.42E-05 | 9.68E-05 | TRUE |
| STY_Substrate | 15135 | 2256 | 7750 | 1313 | 3.04E-13 | 6.08E-12 | TRUE |
| STY_Cofactor | 10124 | 1880 | 5156 | 974 | 0.20604212 | 0.41208424 | FALSE |
| STY_Nucleotides/Derivatives | 5108 | 908 | 1802 | 250 | 0.99999998 | 1 | FALSE |
| STY_Metal Ions | 14051 | 2079 | 5760 | 811 | 0.97833614 | 1 | FALSE |
| S_Active Site | 7784 | 1382 | 3184 | 554 | 0.76152932 | 1 | FALSE |
| S_Substrate | 6434 | 1053 | 3089 | 553 | 0.00077415 | 0.0025805 | TRUE |
| S_Cofactor | 4611 | 906 | 2141 | 379 | 0.99915792 | 1 | FALSE |
| S_Nucleotides/Derivatives | 2237 | 510 | 686 | 99 | 1 | 1 | FALSE |
| S_Metal Ions | 6230 | 1090 | 2290 | 338 | 0.99999482 | 1 | FALSE |
| T_Active Site | 6393 | 722 | 2775 | 349 | 0.00262662 | 0.00656654 | TRUE |
| T_Substrate | 5306 | 570 | 2733 | 326 | 0.0022902 | 0.00654343 | TRUE |
| T_Cofactor | 3520 | 491 | 1850 | 261 | 0.40605114 | 0.66699799 | FALSE |
| T_Nucleotides/Derivatives | 1722 | 173 | 639 | 56 | 0.92649076 | 1 | FALSE |
| T_Metal Ions | 4799 | 487 | 2037 | 213 | 0.28742236 | 0.52258612 | FALSE |
| Y_Active Site | 4054 | 809 | 2018 | 472 | 3.04E-08 | 1.52E-07 | TRUE |
| Y_Substrate | 3395 | 633 | 1928 | 434 | 1.31E-11 | 1.31E-10 | TRUE |
| Y_Cofactor | 1993 | 483 | 1165 | 334 | 2.01E-08 | 1.34E-07 | TRUE |
| Y_Nucleotides/Derivatives | 1149 | 225 | 477 | 95 | 0.43354869 | 0.66699799 | FALSE |
| Y_Metal Ions | 3022 | 502 | 1433 | 260 | 0.01790061 | 0.03977913 | TRUE |

**Table S11:** CRISPRi-rescue GSTP1 activity assay. Associated with. **Figure 6B box and whisker plot representing CDNB day assay on A549 cells treated with doxycycline for 96 hours to measure activity of GSTP1 variants (N=4).**

| SampleID | Replicate 1 | Replicate 2 | Replicate 3 | Replicate 4 |
| --- | --- | --- | --- | --- |
| mCh + Dox | 0.251466852 | 0.240467027 | 0.241174152 | 0.265649952 |
| WT + Dox | 1.139896704 | 1.110555397 | 1.093679865 | 1.17507856 |
| 8F + Dox | 0.381642413 | 0.337951357 | 0.321111623 | 0.325328852 |
| 8E + Dox | 0.352247204 | 0.378501749 | 0.335827965 | 0.330149259 |
| 64F + Dox | 0.783388799 | 0.838407551 | 0.96400229 | 0.86330551 |
| 64E + Dox | 0.363930662 | 0.416270866 | 0.416270866 | 0.365957486 |
| 109F + Dox | 0.652817322 | 0.807335321 | 0.807335321 | 0.785395438 |
| 109E + Dox | 0.417099813 | 0.458795708 | 0.458795708 | 0.400326754 |
| 199F + Dox | 0.886322186 | 0.890084997 | 0.890084997 | 0.809788453 |
| 199E + Dox | 0.874027948 | 0.900584857 | 0.900584857 |  |

**Table S12:** In vitro GSTP1 assay on recombinant protein. Associated with. **Figure 6D CDNB decay assay performed on 1µg of recombinant protein purified from SRC-hyperphosphorylated lysates. mCh=mCherry, WT=wildtype.**

| SampleID | Replicate 1 | Replicate 2 | Replicate 3 | Replicate 4 |
| --- | --- | --- | --- | --- |
| mCh -SRC | 0.01960062 | 0.096603514 | 0.463395879 | 0.005917862 |
| WT -SRC | 8.529107059 | 7.020319251 | 8.646649139 | 6.319226544 |
| 8F -SRC | 0.182939817 | 0.1871399 |  | 0.340813454 |
| 8E -SRC | 0.053201842 | 0.146538308 | 0.913635396 | -0.478083506 |
| 64F -SRC | 7.685343114 | 5.965614842 | 7.946925683 | 3.062577272 |
| 64E -SRC | 0.618355283 | 1.080838987 | 2.230245608 | 0.840197382 |
| 109F -SRC | 7.753012319 | 6.835512669 | 8.767493008 | 4.129983153 |
| 109E -SRC | 1.566189524 | 1.042570712 | 1.280064378 | 1.150241483 |
| 199F -SRC | 7.606007112 | 4.356957029 | 8.311405832 | 3.656631819 |
| 199E -SRC | 6.900848347 | 6.529368208 | 6.598155381 | 2.930036923 |
| mCh +SRC | -0.157272557 | -0.007467201 | 0.402973654 | -0.667424745 |
| WT +SRC | 6.629238979 | 7.017519103 | 6.881748242 | 4.671968983 |
| 8F +SRC | 0.014000463 | 0.202073815 |  | 0.530152224 |
| 8E +SRC | 0.214207791 | 0.204873824 | 0.738217252 | 0.336078812 |
| 64F +SRC | 7.036652823 | 5.577800505 | 7.455755749 | 4.74060348 |
| 64E +SRC | 0.572620363 | 0.455949747 | 1.226464809 | 0.939601938 |
| 109F +SRC | 6.842046487 | 8.195894847 | 6.89831564 | 4.06134795 |
| 109E +SRC | 0.999169179 | 0.597821558 | 2.589852515 | 0.946701256 |
| 199F +SRC | 6.663773027 | 4.381224146 | 5.497894602 | 4.693268347 |
| 199E +SRC | 1.621725032 | 3.176247481 | 4.340135427 | 2.766732865 |

**Table S13:** Data for Figure S6C, western blot and quantitation of CRISPRi-mediated knockdown and rescue of human GSTP1 with a pool of 4 single guide RNAs (sgGSTP1) for 96 hours with 500ng/mL of Dox in A549 cells (N=4).

| **GSTP1_Rep1** | **GSTP1** | **GSTP1_PCNA** | **GSTP1/PCNA** | **FC -Dox/+Dox** |
| --- | --- | --- | --- | --- |
| **sgGSTP1 -Dox** | 61900 | 9920 | 6.239919355 | 100% |
| **sgGSTP1 +Dox** | 27800 | 8670 | 3.206459054 | 51% |

| **GSTP1_Rep2** | **GSTP1** | **GSTP1_PCNA** | **GSTP1/PCNA** | **FC -Dox/+Dox** |
| --- | --- | --- | --- | --- |
| **sgGSTP1 -Dox** | 95700 | 8150 | 11.74233129 | 100% |
| **sgGSTP1 +Dox** | 43700 | 7320 | 5.969945355 | 51% |

| **GSTP1_Rep3** | **GSTP1** | **GSTP1_PCNA** | **GSTP1/PCNA** | **FC -Dox/+Dox** |
| --- | --- | --- | --- | --- |
| **sgGSTP1 -Dox** | 108000 | 34200 | 3.157894737 | 100% |
| **sgGSTP1 +Dox** | 46300 | 31400 | 1.474522293 | 47% |

| **GSTP1_Rep4** | **GSTP1** | **GSTP1_PCNA** | **GSTP1/PCNA** | **FC over -Dox** |
| --- | --- | --- | --- | --- |
| **sgGSTP1 -Dox** | 36400 | 49000 | 0.742857143 | 100% |
| **sgGSTP1 +Dox** | 4160 | 37800 | 0.11005291 | 15% |

**Table S15:**
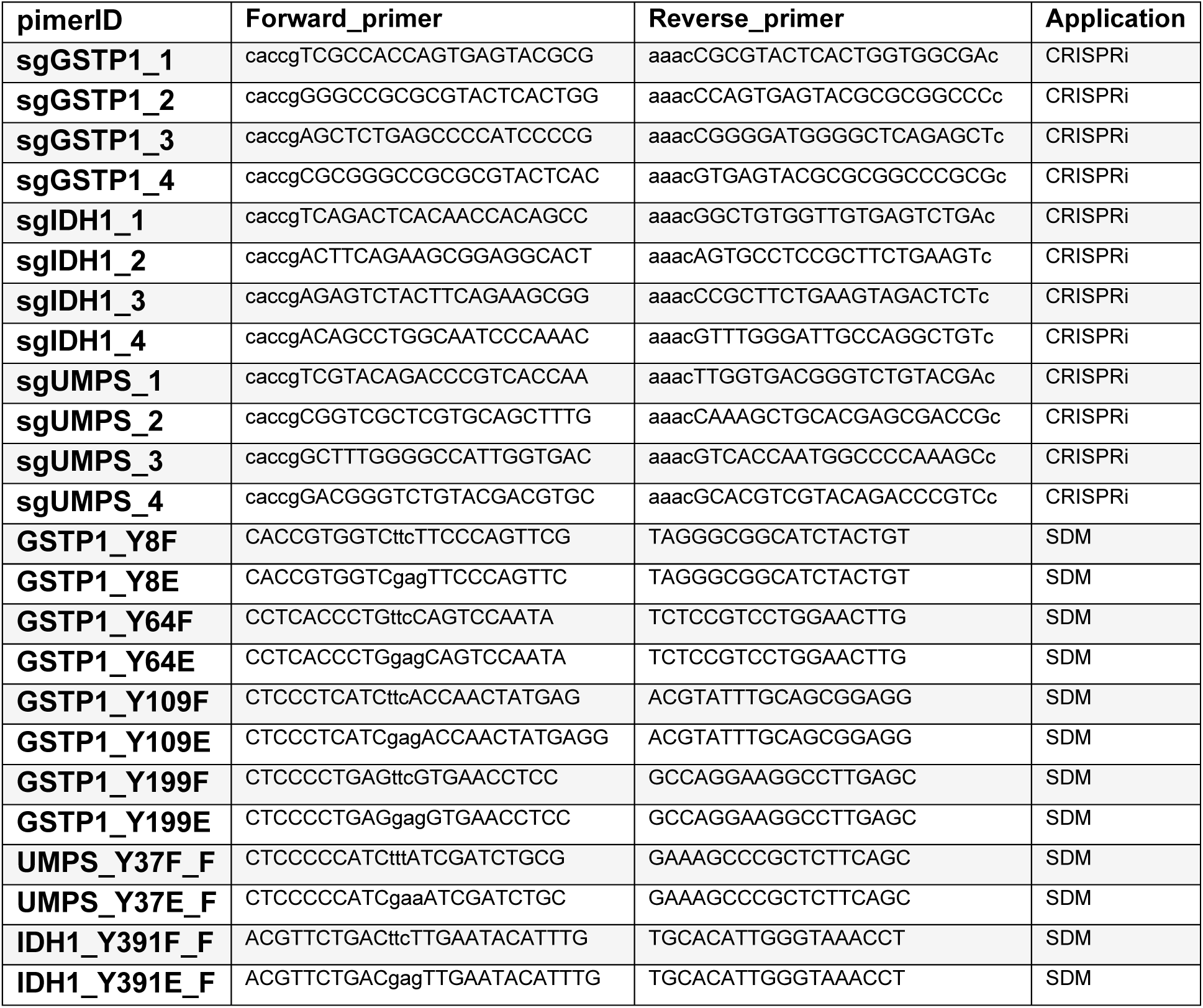
Oligonucleotides used for CRISPRi guides and site-directed mutagenesis.

